# Requirements for the Biogenesis of [2Fe-2S] Proteins in the Human and Yeast Cytosol

**DOI:** 10.1101/2024.01.15.575444

**Authors:** Joseph J. Braymer, Oliver Stehling, Martin Stümpfig, Ralf Rösser, Farah Spantgar, Catharina M. Blinn, Ulrich Mühlenhoff, Antonio J. Pierik, Roland Lill

## Abstract

The biogenesis of iron-sulfur (Fe/S) proteins entails the synthesis and trafficking of Fe/S clusters, followed by their insertion into target apoproteins. In eukaryotes, the multiple steps of biogenesis are accomplished by complex protein machineries in both mitochondria (ISC) and cytosol (CIA). The underlying biochemical pathways have been elucidated over the past decades, yet the mechanisms of cytosolic [2Fe-2S] protein assembly have remained ill-defined. Similarly, the precise site of glutathione (GSH) requirement in cytosolic and nuclear Fe/S protein biogenesis is unclear, as is the molecular role of the GSH-dependent cytosolic monothiol glutaredoxins (cGrxs). Here, we investigated these questions in human and yeast cells by various *in vivo* approaches. [2Fe-2S] cluster assembly of cytosolic target apoproteins required the mitochondrial ISC machinery, the ABC transporter Atm1/ABCB7 and GSH, yet occurred independently of both the CIA system and cGrxs. This mechanism was strikingly different from the ISC-, Atm1/ABCB7-, GSH-, and CIA-dependent assembly of cytosolic-nuclear [4Fe-4S] proteins. One notable exception to this newly defined cytosolic [2Fe-2S] protein maturation pathway was the yeast protein Apd1 which used the CIA system via binding to the CIA targeting complex through its C-terminal tryptophan. cGrxs, although attributed as [2Fe-2S] cluster chaperones or trafficking proteins, were not essential *in vivo* for delivering [2Fe-2S] clusters to either CIA components or apoproteins. Finally, GSH function was assigned to Atm1-dependent export, i.e. a step before GSH-dependent cGrxs function. Our findings extend the general model of eukaryotic Fe/S protein biogenesis by adding the molecular requirements for cytosolic [2Fe-2S] protein maturation.

## Introduction

Iron-sulfur (Fe/S) clusters are synthesized and inserted into eukaryotic target apoproteins by evolutionary conserved protein machineries in mitochondria and cytosol (1–3). In mitochond r ia, the iron-sulfur cluster assembly (ISC) system is responsible for generating [2Fe-2S] and [4Fe-4S] proteins within the organelle (Fig. S1). Early-acting ISC components comprise the cysteine desulfurase complex Nfs1-Isd11-Acp1 (yeast nomenclature, refer to Table S1 for human protein names), the electron donor ferredoxin Yah1, and the allosteric effector Yfh1 (yeast frataxin homolog), which together *de novo* synthesize a [2Fe-2S] cluster on the scaffold protein Isu1 (4). The cluster is released from Isu1 by the dedicated Hsp70/Hsp40 chaperones Ssq1 and Jac1, transiently trafficked to monothiol glutaredoxin Grx5, and either inserted into target [2Fe-2S] apoproteins or used by late-acting ISC factors for Yah1-dependent fusion to [4Fe-4S] clusters and insertion into apoproteins (5, 6). Additionally, the early ISC components produce a still unknown sulfur- and iron-containing substrate (X-S) that is exported by the mitochondr ia l ABC transporter Atm1 to support the cytosolic iron-sulfur protein assembly (CIA) machinery (Fig. S1) (4, 7–9).

The CIA system initiates the synthesis of a [4Fe-4S] cluster on the cytosolic scaffold complex Cfd1-Nbp35, a reaction depending on the electron transport chain NADPH-Tah18-Dre2 (1, 10). The [4Fe-4S] cluster is then presumably passed on via Nar1 to the CIA targeting complex (CTC) composed of Cia1-Cia2-Mms19 that mediate the maturation of target [4Fe-4S] proteins like DNA maintenance enzymes (11–14).

Maturation of cytosolic and nuclear [4Fe-4S] proteins also requires the tripeptide glutathione (GSH) (15, 16), yet the exact site of GSH requirement is not clear. Since GSH has been shown to bind to Atm1 (8), and since GSH-containing compounds such as GSSSG and a tetra-GSH- coordinated [2Fe-2S] cluster have been suggested as potential candidates for the enigmatic X-S substrate (9, 17), GSH may be directly connected to the Atm1-dependent export process. Alternatively, the GSH requirement may be connected to GSH-binding of cytosolic monothiol glutaredoxins (cGrxs; Grx3-Grx4) which are important for cytosolic-nuclear [4Fe-4S] protein assembly and also cellular iron regulation (18–21). Eukaryotic cGrxs are composed of an N-terminal thioredoxin domain and up to three C-terminal monothiol glutaredoxin (Grx) domains. *cGRX* gene deletion in various eukaryotes is associated with a surprisingly different severity of phenotypes, from being lethal to hardly any consequences for viability (Table S2). Yeast Grx3- Grx4, in complex with the cytosolic BOLA family protein Bol2, are essential for Aft1-Aft2- dependent transcriptional iron regulation (19–23). Further, cGrxs and their complex with BOLA2 have been linked to cellular iron homeostasis in animals, and were suggested to act as [2Fe-2S] cluster chaperones or trafficking factors for the maturation of human CIAPIN1 and NBP35 (24–28).

Contrary to the advances in understanding how the CIA system synthesizes, traffics, and inserts [4Fe-4S] clusters of target apoproteins (1, 13), the *in vivo* process of cytosolic [2Fe-2S] protein biogenesis has remained poorly characterized (Fig. S1). Relatively few cytosolic [2Fe-2S] proteins per species are known to date (Fig. S1) (4, 19, 29–33). In addition to true target proteins, also the cGrxs and the early CIA components Dre2, Cfd1, and Nbp35 have been reported to (transiently) bind [2Fe-2S] clusters *in vitro* (and *in cellulo* for Dre2) suggesting that also these proteins are acceptors of binuclear clusters (26, 34–37). Currently, it is unclear if and to what extent the CIA system may play a role in cytosolic [2Fe-2S] protein maturation. While some *in vivo* studies have indicated that a functional CIA system is dispensable for [2Fe-2S] cluster assembly of cytosolic [2Fe-2S] target proteins (29, 38), other *in vivo* and *in vitro* studies have suggested a CIA dependence (39–44). A major goal of the current study was to clarify this apparent discrepancy, and to define the *in vivo* trafficking and insertion pathways for cytosolic [2Fe-2S] proteins using yeast and human cells as models. Further, we explored the most critical site of GSH requirement in cytosolic-nuclear Fe/S protein biogenesis, and we investigated the involvement of cGrxs in cytosolic Fe/S protein biogenesis. Together, our results complement and expand the current view of cytosolic Fe/S protein biogenesis in eukaryotes.

## Results

### Human cytosolic [2Fe-2S] protein maturation requires the ISC but not the CIA system

To begin defining the requirements for cytosolic [2Fe-2S] protein assembly, we first tested its dependence on ISC and CIA components in cultured human cells. Aldehyde oxidase contains both [2Fe-2S] clusters and molybdenum cofactors (MoCo) (45), yet the insertion of the Fe/S cluster occurs before and independently of MoCo (46). In order to probe Fe/S cluster binding, we employed a ^55^Fe radiolabeling-immunoprecipitation assay (Supplemental Methods and (47, 48)). Murine aldehyde oxidase (Aox, fused to an N-terminal myc-TEV-EGFP tag) was expressed either transiently from a plasmid in HeLa cells or was inducibly produced in HEK293 FlpIn T-REx cells, each depleted for various ISC and CIA components by RNAi and cultured in presence of ^55^Fe (Fig. 1A and S2). At cell harvest, total protein yield from the differe nt depletion conditions varied only moderately, and the myc-TEV-EGFP-Aox reporter construct did not influence the ^55^Fe content in cell extracts (Fig. S2A-C). Upon RNAi-mediated depletion of the ISC protein NFS1 or the export component ABCB7, we observed a 50% decrease in Aox-associated ^55^Fe, as compared to the corresponding control cells (Fig. 1A and S2A,D). Complementation with RNAi-resistant versions of *NFS1* or *ABCB7* recovered ^55^Fe incorporation into Aox substantially. In contrast, depletion of the CIA proteins NBP35 (NUBP1) or CIAO3 (yeast Nar1) did not decrease ^55^Fe binding to Aox. Complementation with RNAi-resistant constructs was without major effects. Moreover, individual or combined depletion of the CIA components CIAPIN1 and NDOR1 (yeast Dre2 and Tah18) elicited even a twofold increase in ^55^Fe incorporation into Aox (Fig. 1A and Fig. S2A,D). Taken together, [2Fe-2S] cluster maturation of Aox depends on the mitochondrial ISC system and ABCB7- dependent export, yet is not hampered or even increased upon depletion of CIA factors.

**Figure 1.**
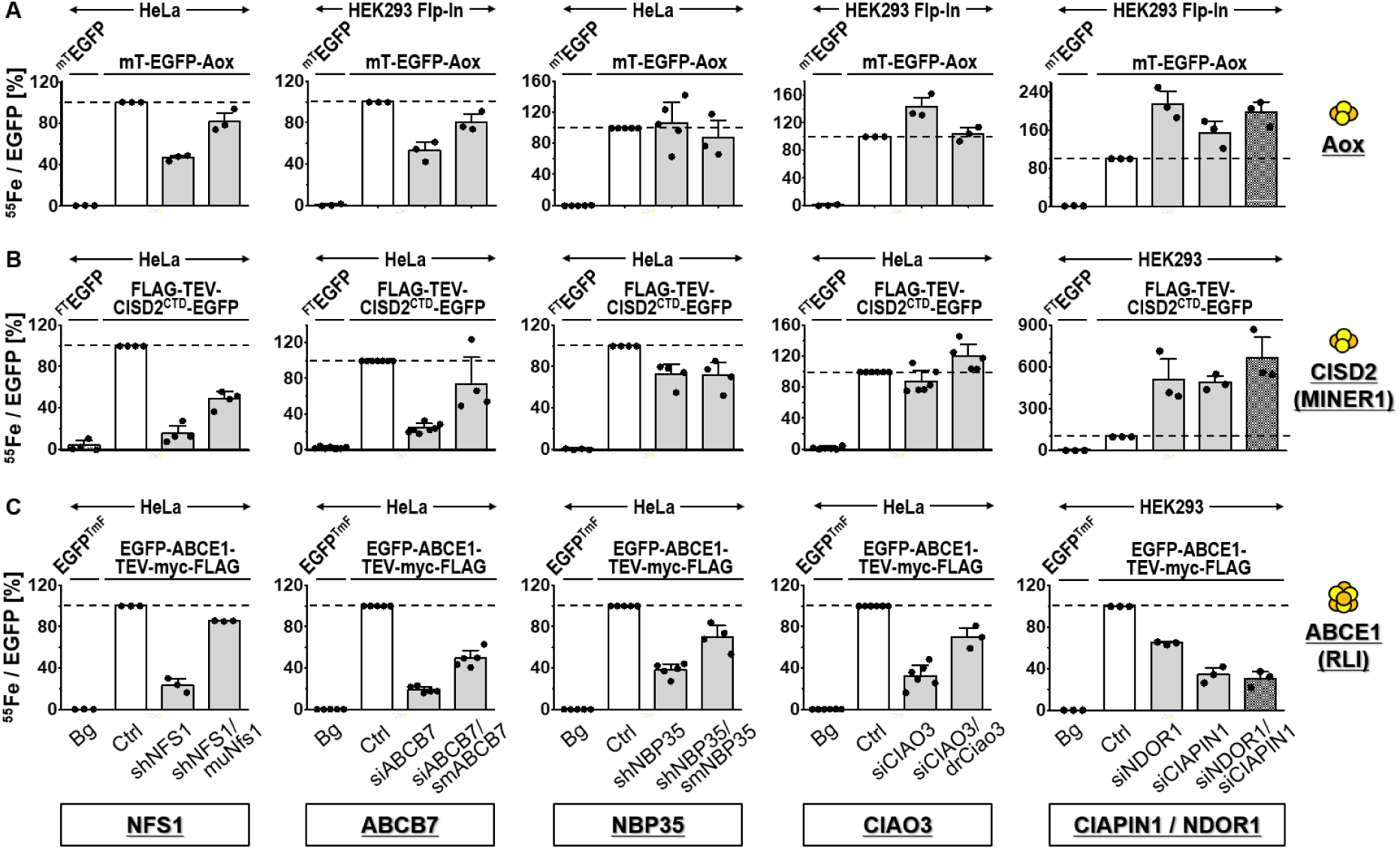
Human cytosolic [2Fe-2S] protein maturation requires ISC but not CIA components. HeLa, HEK293, or HEK293 Flp-In cells were individually or double-depleted for NFS1, ABCB7, NBP35, CIAO3, NDOR1, or CIAPIN1 for a total of six days by transfection with shRNA or siRNA. In parallel, RNAi-treated cells were complemented by RNAi-resista nt murine (mu) NFS1, silently mutated (sm) ABCB7, smNBP35, or *Danio rerio* (dr) Ciao3 as indicated. Control (Ctrl) samples did not receive siRNA or shRNA. We concomitant ly expressed the [2Fe-2S] reporter proteins (**A**) myc-TEV-EGFP-Aox and (**B**) FLAG-TEV-CISD2^CTD^-EGFP, or (**C**) the [4Fe-4S] EGFP-ABCE1-TEV-myc-FLAG, either transiently for three days (HeLa, HEK293) or inducibly by doxycycline treatment (HEK293 Flp-In) for 18 h prior to harvest. Myc- and FLAG-tagged EGFP contructs served as reference for non-specific ^55^Fe background binding (Bg, first bars). After ^55^Fe radiolabeling for three days, maturation of reporter proteins was assessed by α-FLAG or α-myc immunoprecipitation, EGFP analysis, and scintillation counting of the recovered material. Fe/S cluster assembly is expressed as the ratio of recovered ^55^Fe radioactivity per EGFP fluorescence, and values (mean ± SD, n≥3) are normalized to those obtained for reporter-expressing control cells (set to 100%, white bars and dashed lines). Abbreviations: sh, small hairpin; si, small interfering; CTD, C-terminal domain; mT, myc-TEV; FT, FLAG-TEV; TmF, TEV-myc-FLAG.

We further tested CISD2 (aka MINER1, NAF-1), a cytosolic member of the [2Fe-2S] cluster-binding NEET family containing a 3Cys-1His coordination motif (49). The soluble C-terminal domain (CTD) of CISD2 was employed to construct a fusion protein with an N-terminal FLAG-TEV tag and a C-terminal EGFP. This cytosolic FLAG-TEV-CISD2(CTD)-EGFP reporter was expressed in HeLa or HEK293 cells in parallel to the depletion of ISC, ABCB7, or CIA proteins. Similar to Aox, ^55^Fe binding to CISD2(CTD) was dependent on functional NFS1 and ABCB7, yet did not require NBP35 or CIAO3 (Fig. 1B and Fig. S3). Individual or combined deficiency of CIAPIN1 and NDOR1 increased the ^55^Fe incorporation several-fold, despite unchanged cellular ^55^Fe levels (Fig. S3C). As a control, we studied the maturation of the [4Fe-4S] protein ABCE1 (11, 47, 50) under the conditions used above. As expected, ^55^Fe binding to ABCE1 was specifically dependent on mitochondrial ISC, ABCB7 and CIA proteins (Fig. 1C, Fig. S4). Collectively, our results from human cell culture, together with previous studies on CISD1 (38), clearly indicate the mitochondria-dependent, yet CIA-independent maturation of cytosolic [2Fe-2S] proteins, distinguishing this pathway from that of cytosolic [4Fe-4S] protein assembly.

### CIAPIN1 is matured independently of CIA and GLRX3 mimicking [2Fe-2S] target proteins

CIAPIN1 and its relatives contain at least one (yeast Dre2; (35, 36, 51)) or two (human CIAPIN1; (37, 52)) [2Fe-2S] clusters. *In vitro* and *in vivo* studies have suggested that human CIAPIN1 may receive its [2Fe-2S] clusters by transfer from GLRX3 (27) or a GLRX3-BOLA2 heterocomplex (25). In order to study the requirements for CIAPIN1 maturation in human cells, we applied a CRISPR approach to fuse endogenous CIAPIN1 to a genomically integrated C-terminal EGFP-TEV-myc-FLAG tag. Using the ^55^Fe radiolabeling-immunoprecipitation assay (cf. Fig. 1), maturation of the tagged CIAPIN1 was probed upon RNAi-mediated depletion of the core ISC component ISCU2 or of the cytosolic proteins NDOR1, GLRX3, or NBP35 (Fig. 2A). Gene silencing was efficient (Fig. S5A-B), and resulted in a slight growth defect as measured by protein content (Fig. S5C), yet no effects on cellular iron uptake (Fig. S5D). Substantial amounts of ^55^Fe were co-immunoprecipitated with CIAPIN1-EGFP-TEV-myc-FLAG, while cells expressing a myc-TEV-EGFP reference construct bound only background levels of ^55^Fe (Bg, Fig. 2A, Fig. S5E). Depletion of ISCU2 resulted in a 50% decrease of CIAPIN1-associated ^55^Fe, consistent with the dependency of all cytosolic-nuclear Fe/S proteins on the mitochondrial ISC system (cf. Fig. 1). In contrast, knock-down of the CIA proteins NDOR1 and NBP35 did not diminish but rather increased CIAPIN1-bound ^55^Fe (Fig. 2A, Fig. S5E). These results fit well to the independence of cytosolic [2Fe-2S] protein maturation on the CIA factors (see above).

**Figure 2.**
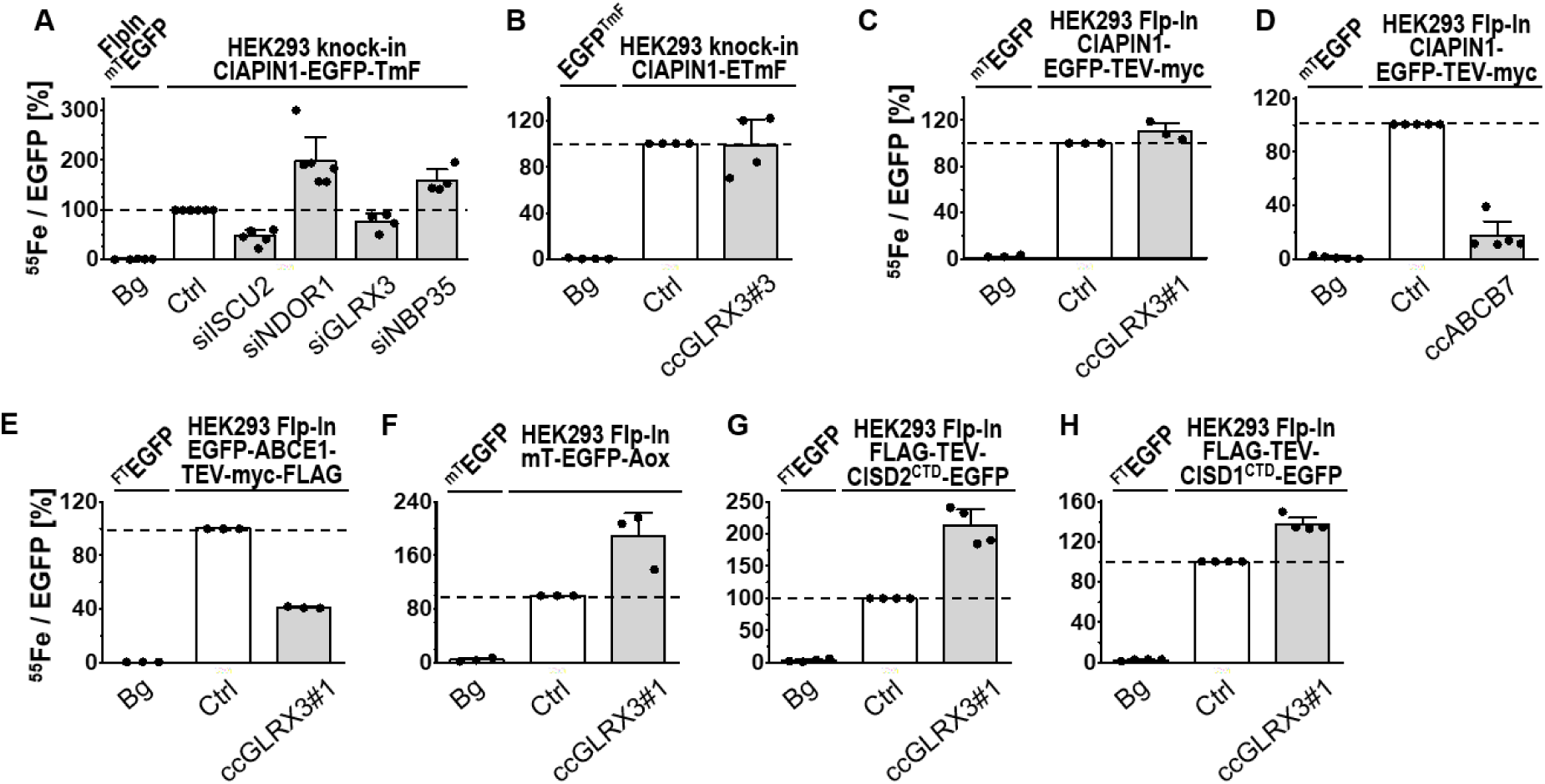
Maturation of human [2Fe-2S]-containing CIAPIN1 occurs independently of GLRX3. **(A)** HEK293 cells expressing endogenous CIAPIN1 fused to a genome-integrated C- terminal tag encoding EGFP-TEV-myc-FLAG (knock-in) were depleted for the indicated proteins by siRNA treatment for 6 days. Radiolabeling with ^55^Fe and maturation of the CIAPIN1 fusion protein was analyzed in comparison to control-transfected cells (Ctrl) according to Fig. 1. **(B)** *GLRX3* was knocked out in untreated CIAPIN1 knock-in cells from part (A) by CRISPR-Cas9 (construct #3; cf. Fig. S6B), and ^55^Fe/S cluster maturation was analyzed as in (A) relative to control-transfected cells. (**C-D**) *GLRX3* (construct #1; cf. Fig. S6B) or *ABCB7* genes were knocked out by CRISPR-Cas9 in HEK293 Flp-In cells inducib ly expressing CIAPIN1-EGFP-TEV-myc. ^55^Fe/S cluster maturation was analyzed as in (B). **(E-H)** *GLRX3* knock-out cells from (C) inducibly expressing the indicated Fe/S reporter proteins were analyzed for ^55^Fe/S cluster maturation as in Fig. 1. Cells with inducibly or transient ly expressed myc- or FLAG-tagged EGFP variants served as ^55^Fe background binding reference (Bg). Values are presented relative to the reporter protein-expressing Ctrl cells (set to 100%, white bars and dashed lines) and are given as mean ± SD, n ≥ 3. For abbreviations see Fig. 1.

Surprisingly, GLRX3 depletion by a validated siRNA (24) also did not critically affect ^55^Fe incorporation into CIAPIN1 (Fig. 2A, Fig. S6E), contrary to earlier findings (25). Since residual GLRX3 levels, due to incomplete siRNA-mediated depletion, might be sufficient to support CIAPIN1 maturation, we used a CRISPR/Cas9 approach (53) to knock out the *GLRX3* gene in HEK293 cells. Transfection of puromycin resistance cassette-containing plasmids encoding three different *GLRX3*-directed gRNAs (ccGLRX3#1-3) yielded viable cell lines after antibiotics selection (Fig. S6A). All three gRNAs were highly efficient in depleting GLRX3 (Fig. S6B), but none of the gRNA-treated cells was severely growth-retarded (cf. Fig. S6A, indicating that GLRX3 is not essential for viability of cultured human cells. This behavior is similar to the *cGRX* gene knockout in plants and some fungi, yet clearly different for other fungi or mice, where *cGRX* deletion is lethal (Table S2). Application of the ^55^Fe radiolabeli ng-immunoprecipitation assay to the *GLRX3* knock-out cells revealed an unchanged cellular ^55^Fe content (Fig. S6C) and inconspicuous ^55^Fe maturation of genome-integrated CIAPIN1-EGFP-TEV-myc-FLAG (Fig. 2B, Fig. S6C-E), thus verifying the RNAi depletion results above.

In a second approach, we inducibly overproduced CIAPIN1-EGFP-TEV-myc in HEK293 FlpIn T-Rex cells, but again did not observe any effect of a CRISPR-mediated *GLRX3* knockout (KO) on ^55^Fe incorporation into CIAPIN1 (Fig. 2C). The cells contained normal levels of endogenous CIAPIN1, and showed regular cell growth and cellular ^55^Fe levels (Fig. S7A-D). In contrast to GLRX3, CRISPR-Cas9-mediated knock-out of *ABCB7* elicited a severe defect in ^55^Fe binding by inducibly overproduced CIAPIN1 (Fig. 2D, Fig. S7E,H), consistent with the general function of ABCB7 in cytosolic-nuclear Fe/S protein assembly. The *ABCB7*-depleted cells grew rather poorly after puromycin selection, but maintained an almost regular cellular ^55^Fe content (Fig. S7F,G). Collectively, maturation of the [2Fe-2S] CIA protein CIAPIN1 depends on mitochondrial ISC but not on CIA and GLRX3.

### GLRX3 knockout affects assembly of cytosolic [4Fe-4S] but not [2Fe-2S] proteins

The independence of human CIAPIN1 on GLRX3 raised the question of the role of the latter in cytosolic [2Fe-2S] and [4Fe-4S] protein assembly. As reported earlier (18) and in this work (see below), yeast Grx3-Grx4 play a role in cytosolic [4Fe-4S] but not [2Fe-2S] protein maturation. Similarly, previous work found that *GLRX3*-depleted HeLa cells showed twofold decreases of cytosolic aconitase (IRP1) activity and GPAT protein levels, again linking human GLRX3 to [4Fe-4S] protein maturation (24). This connection was further generalized by ^55^Fe radiolabeling of our *GLRX3* KO HEK293 cells. We observed a 60% decrease of ^55^Fe binding to the reporter EGFP-ABCE1-TEV-myc-FLAG in the absence of GLRX3 (Fig. 2E; Fig. S8 left most panels). This result indicated an important but not essential function of GLRX3 in ABCE1 maturation. Notably, ABCE1 and numerous other [4Fe-4S] proteins including DNA polymerases and helicases (54) are essential for cell viability. Nevertheless GLRX3-deficient cells grew well (cf. Figs. S6C and 7A), further verifying that in cultured cells GLRX3 function is dispensable.

For the analysis of cytosolic [2Fe-2S] protein maturation, we inducibly expressed myc-TEV-EGFP-Aox, FLAG-TEV-CISD2(CTD)-EGFP, and FLAG-TEV-CISD1(CTD)-EGFP in *GLRX3* knockout cells. ^55^Fe radiolabeling did not reveal a requirement of GLRX3 for their maturation (Fig. 2F-H, Fig. S8D, right panels). Rather, the GLRX3 deficiency led to an up to twofold higher ^55^Fe association with these [2Fe-2S] proteins compared to control cells, despite of wild-type levels of cellular ^55^Fe(Fig. S8C, right panels). Together, these results show a (non-essential) *in vivo* function of GLRX3 in cytosolic [4Fe-4S] but not [2Fe-2S] protein assembly in cultured human cells.

### Yeast cGrxs impact but are not essential for the CIA pathway

The contrasting cGrxs dependencies of human CIAPIN1 and yeast Dre2 maturation prompted us to more deeply investigate the function of the yeast cGrxs for which a crucial (BY strain background; (55, 56)) or essential (W303; (18)) role in cytosolic-nuclear Fe/S protein biogenesis has been reported (Table S2). Extending earlier studies, Dre2 maturation *in vivo* required Grx3-Grx4 in addition to the mitochondrial ISC system (18, 34), but not the CIA machinery, unlike Leu1 (Fig. S9). Depletion of the cGrx proteins also results in a massive accumulation of iron in the cell, which for unknown reasons cannot be properly incorporated into iron-containing proteins (18). In order to tease apart whether this pool of non-bioavailable iron impacts cytosolic Fe/S protein biogenesis, the gene of the major transcriptional regulator of the iron uptake system, Aft1, was deleted in addition to depleting the cGrxs (yielding Gal-*GRX4grx3Δaft1Δ* cells (*cGRX↓aft1Δ*)). Depletion of Grx4 in this regulatable yeast strain was achieved by the replacement of the natural promoter with the glucose-repressing and galactose-inducing *GAL* promoter (depletion is indicated by ↓ in a **l** Figures, for depletion conditions see Table S3) (57). Strikingly, the lethal phenotype of cGrxs in *S. cerevisiae* W303 strains was alleviated upon *AFT1* deletion and *cGRX↓aft1Δ* cells showed almost wild-type growth (Fig. 3A, Table S2). As expected, *AFT1* deletion suppressed the expression of a GFP reporter protein from the Aft1-dependent *FET3* promoter (Fig. 3B) (58). Effects on Fe/S protein maturation in this yeast strain were further studied by *in vivo* ^55^Fe radiolabeling-immunoprecipitation (59). As expected (60), *AFT1* deletion returned cellular iron levels to WT levels (Fig. 3C). Despite the rescue of growth of *cGRX↓aft1Δ* cells, ^55^Fe binding to immunoprecipitated endogenous Dre2 or Leu1 remained as low as in *cGRX↓* cells (Fig. 3D-F). Similarly, a strong decrease in ^55^Fe radiolabeling of Dre2 was also observed in *ATM1↓* cells, which cannot supply X-S for cytosolic Fe/S protein assembly. The diminished ^55^Fe levels in Dre2 and Leu1 in *cGRX↓aft1Δ* cells were cGrx-related, as the deletion of *AFT1* alone did not affect ^55^Fe binding to either of the proteins. These results indicate that the lack of yeast cGrxs impairs Dre2 maturation even when deleterious iron is removed in *cGRX↓aft1Δ* cells, yet the growth assay indicates that Dre2 retains enough residual activity to support cell growth and CIA function.

**Figure 3.**
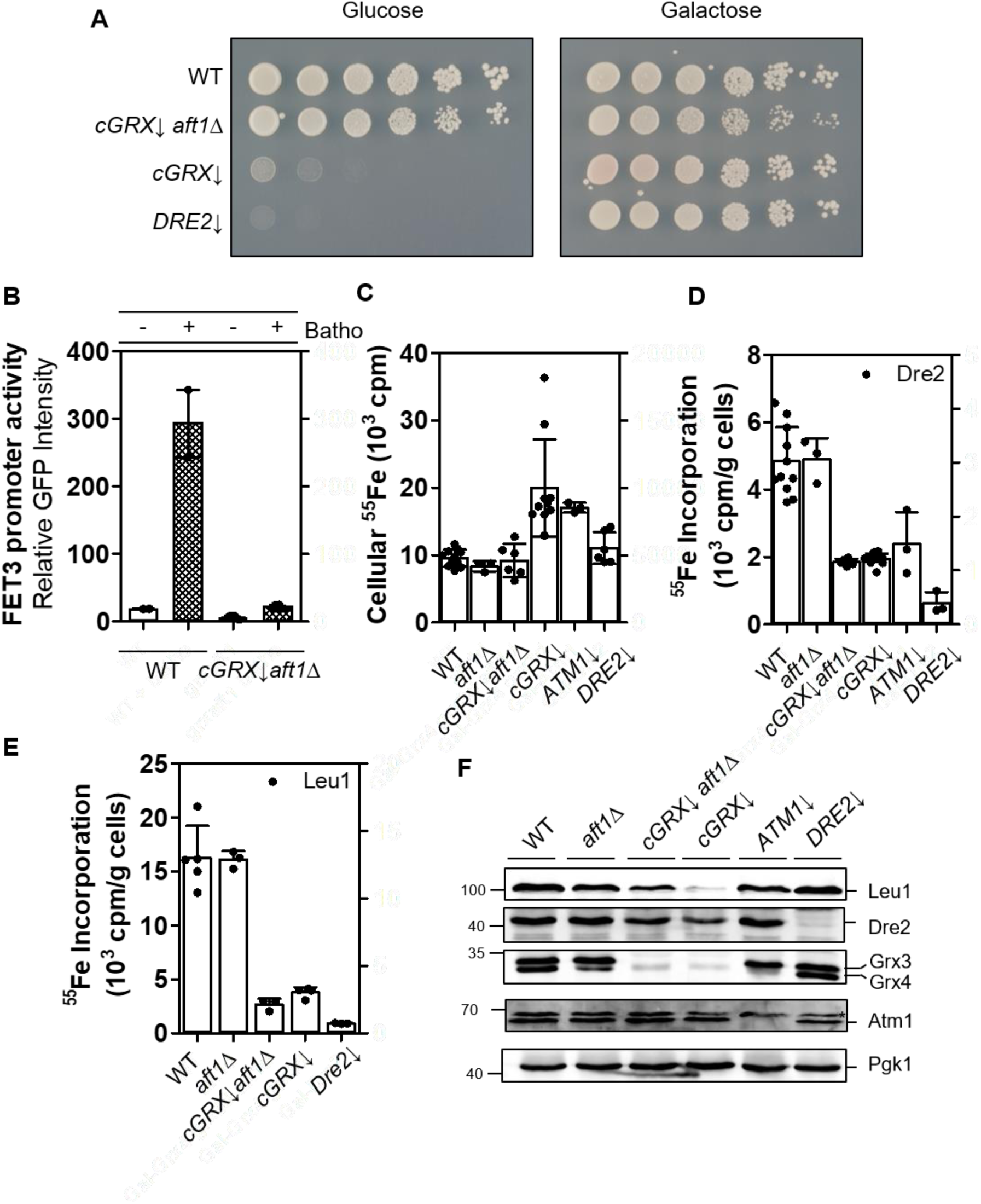
The essential function of cGrxs in *S. cerevisiae* is tied to a combination of Fe/S protein biogenesis and iron regulation. (**A**) The indicated *GAL* promoter-exchange yeast cells (protein depletion indicated by (↓); see Table S3) and wild-type (WT) cells were grown in SC media containing glucose. Serial dilutions (1:5) were plated on glucose- or galactose-containing SC agar plates, and incubated for 3 days at 30°C. (**B**) WT and *cGRX↓aft1Δ* cells were transformed with plasmid pFET3-*GFP,* and grown to mid-log phase with either 50 μM FAC (-) or 50 μM bathophenanthroline (Batho) in glucose-containing SC medium. *FET3* promoter activity was determined by measuring the GFP fluorescence at 513 nm. (**C-E**) WT and the indicated (protein-depleted) yeast strains were subjected to radiolabeling with ^55^FeCl3 for 2 h followed by cell lysis. Using scintillation counting, the ^55^Fe cellular uptake (C) was determined in addition to the amount of ^55^Fe bound to immunoprecipitated Dre2 (D) and Leu1 (E) using homemade antibodies. (**F**) Representative Western blots depict protein levels of Fe/S protein biogenesis components and the loading control Pgk1. Values in bar charts represent the mean ± SD, n ≥ 3 (except for WT samples in B, where n=2, see (63)). *, denotes a non-specific band.

We next focused on the proposed [2Fe-2S] cluster-trafficking function of cGrxs to the scaffold components Cfd1 and Nbp35. As noted above, such transfer has been observed *in vitro* from human GLRX3 to NBP35 (10, 26). cGrx-depleted yeast cells expressing TAP-tagged Cfd1 and Nbp35 either alone or together were radiolabeled with ^55^Fe. TAP affinity-precipitated Nbp35 and/or Cfd1 bound up to three-fold more ^55^Fe (Fig. 4A,B, Fig. S10). In contrast, yet consistent with previous results (18), maturation of the [4Fe-4S] protein Rad3 was strongly dependent on cGrxs. Whereas ^55^Febinding to co-expressed Cfd1-Nbp35 was independent of cGrxs (Fig. 4A), the reaction strongly relied on Atm1 and Tah18-Dre2 functions showing that the bound ^55^Fe is part of a Fe/S cluster (Fig. 4C-E). We conclude that Fe/S cluster binding to yeast Cfd1-Nbp35 does not require cGrxs, making a cluster-trafficking function of cGrxs to Cfd1-Nbp35 unlikely. Moreover, the residual maturation of Dre2 in cGrx-depleted cells (Fig. 3D) was still sufficient to support the crucial function of Dre2 in Fe/S cluster assembly on Cfd1-Nbp35. Overall this indicates that, similarly to the human system, the function of yeast cGrxs in the essential CIA pathway can be bypassed.

**Figure 4.**
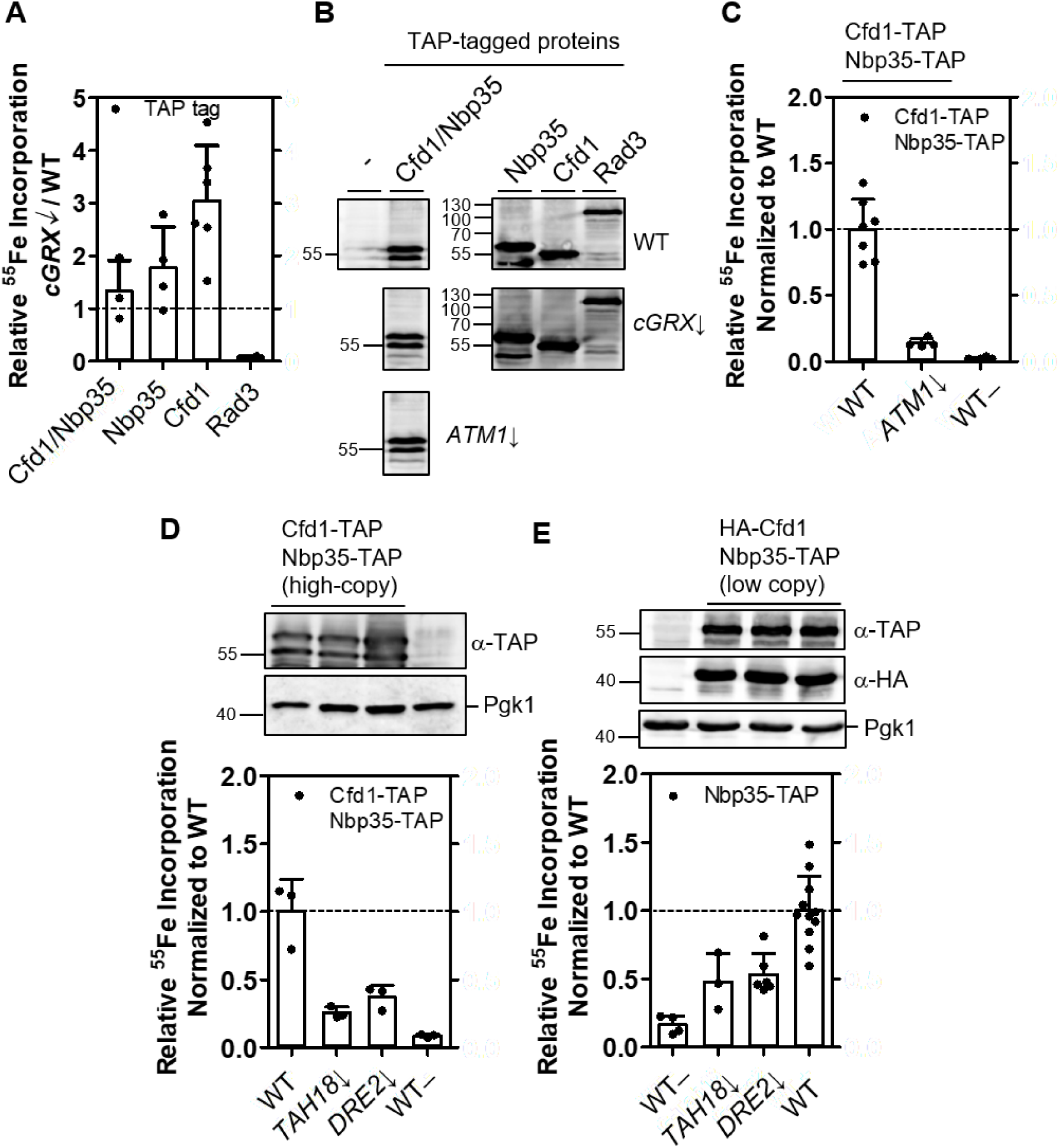
Binding of ^55^Fe to the Cfd1-Nbp35 scaffold complex is independent of cGrxs, but strongly depends on the Atm1-mediated export and Tah18-Dre2. (**A**) Wild-type (WT) and depleted *cGRX*↓ cells expressing TAP-tagged Cfd1 and/or Nbp35 and Rad3 were radiolabeled with ^55^Fe as in Fig. 3C-E. After TAP affinity purification and scintillation counting, protein-bound ^55^Fe was determined. (**B**) Expression of TAP-tagged proteins in WT (top), *cGRX*↓ (middle), and *ATM1*↓ (bottom) strains was estimated by Western blotting. All blots were stained against the TAP tag. (**C-E**) WT cells and strains depleted of *ATM1* (C) or *TAH18* and *DRE2* (D-E) expressing tagged Cfd1-Nbp35 from either high- (C-D) or low-copy (E) vectors were subjected to the ^55^Fe radiolabeling assay as in (A). ^55^Fe binding to Cfd1-Nbp35 is presented relative to that in WT cells. Cells transformed with yeast plasmids lacking epitope tags served as controls (WT-). The top panels in D and E show immunostainings of the indicated proteins, in the order of the bar charts below. Horizontal dashed lines in bar charts represent WT levels of ^55^Fe binding. Values represent the mean ± SD, n ≥ 3, except for Rad3-TAP in (A) (n=2). Uncropped immunostain for the left part of (B) is shown in Fig. S10.

### Yeast cytosolic [2Fe-2S] protein maturation can be CIA-dependent or -independent

As the yeast CIA components Dre2, Nbp35, and Cfd1 could be matured *in vivo* independently of the cGrxs, we assessed in more detail [2Fe-2S] cluster maturation of other yeast proteins. Among the few known yeast cytosolic [2Fe-2S] target proteins are the transcription factors Aft1-Aft2 and Yap5 involved in cellular iron regulation (Fig. S1) (19, 29). For Aft1/2, we could not co-immunoprecipitate any ^55^Fe, even after Aft1 overexpression. This prevented any *in vivo* conclusions on the specific requirements for [2Fe-2S] cluster binding, yet the ISC, Atm1, and cGrx dependence, and CIA independence of the Aft1/2-dependent induction of the iron regulon suggests that the CIA pathway does not influence [2Fe-2S] cluster assembly at Aft1-Aft2 (61–63). For Yap5, we extended the previous *in vivo* analysis (29), and show here that ^55^Fe radiolabeling of the myc-tagged actuator domain responsible for Fe/S cluster binding was independent of the early CIA proteins Tah18, Dre2, and Nbp35 (Fig. S11). Further, Western blotting indirectly confirmed that Yap5 maturation was CIA-independent. In cells depleted of the early ISC proteins, Grx4 protein levels were nearly abolished compared to WT cells (Fig. S12A) (18), because Yap5 regulates Grx4 expression in a [2Fe-2S] cluster-dependent manner (29). In CIA-depleted cells, however, Grx4 levels were maintained or even elevated indicat ing that CIA dysfunction did not interfere with [2Fe-2S] cluster assembly and activation of Yap5 (Fig. S12A). Rather, CIA depletion promoted [2Fe-2S] cluster binding to Yap5 as seen by ^55^Fe radiolabeling (Fig. S11). We conclude that [2Fe-2S] cluster assembly to Aft1-Aft2 and Yap5 is CIA-independent.

We next studied the cytosolic electron transfer protein of unknown function, Apd1, which binds a [2Fe-2S] cluster in a C-terminal thioredoxin-like ferredoxin domain (FD2) (40, 64). The protein structure of Apd1 modelled by AlphaFold2 supports the assignment of [2Fe-2S] cluster binding by showing a 2Cys2His binding pocket in the FD2 domain of Apd1 nearly perfectly aligning with the 4Cys [2Fe-2S] cluster-binding site of a bacterial ferredoxin from *Azotobacter vinelandii* (PDB 5ABR, Fig. 5A-B) (65). Interestingly, recent EPR analysis of overexpressed Apd1 and a functional test of Apd1 (see also below) has suggested that this protein may not follow the CIA independence of cytosolic [2Fe-2S] proteins observed here for yeast and human cells (39, 40). Here, we employed the ^55^Fe radiolabeling-immunoprecipitation assay in order to *in vivo* evaluate the Apd1 maturation dependence on various ISC and CIA proteins.

**Figure 5.**
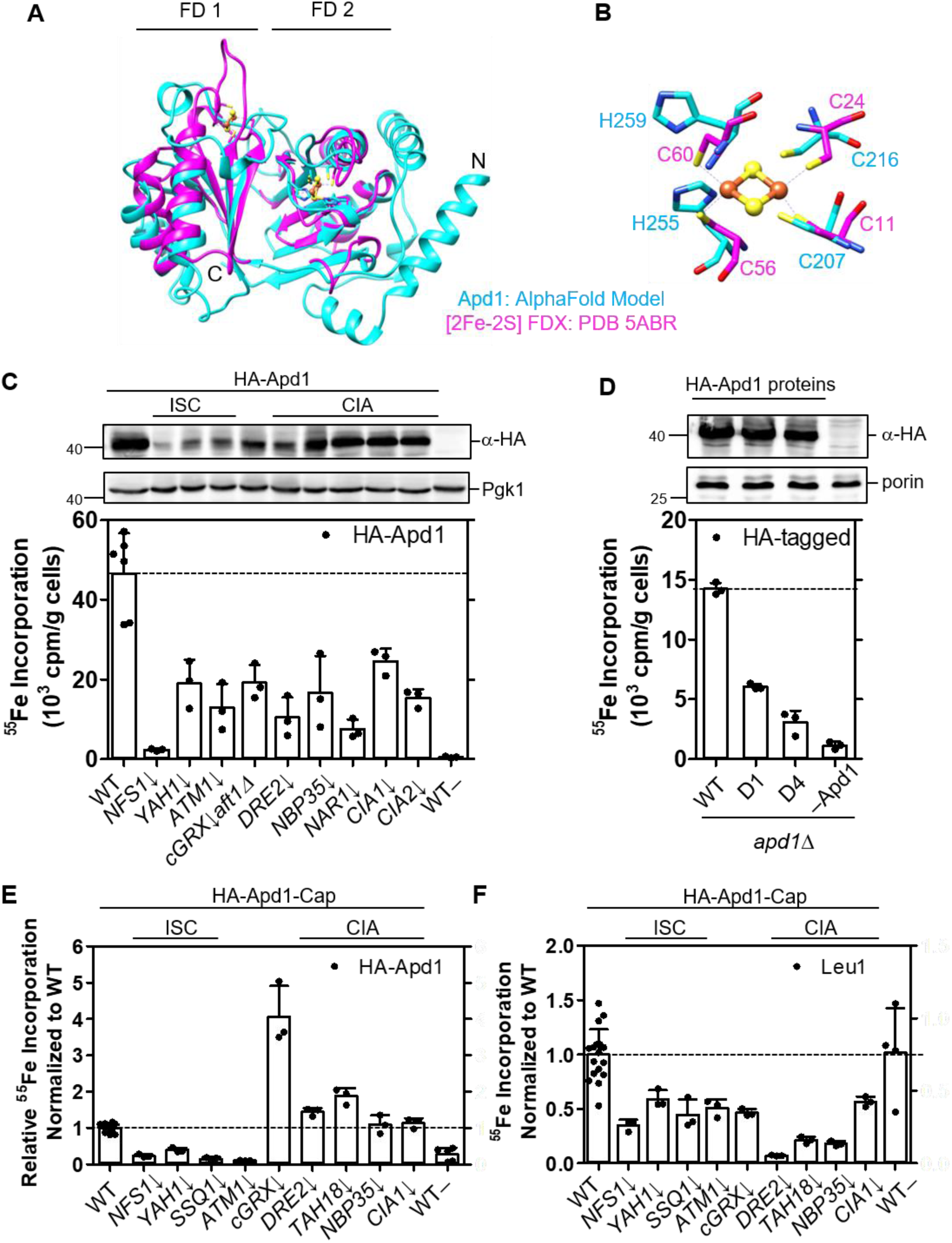
Apd1 shows both an ISC and CIA dependency. (**A**) The AlphaFold-predicted protein structure of Apd1 with two ferredoxin-like domains (FD1 and FD2) is shown in cyan, overlaid with the crystal structure of the dimeric bacterial ferredoxin (in magenta) from *Azotobacter vinelandii* with two [2Fe-2S] clusters (yellow, sulfur; orange, iron). (**B**) The 4Cys coordinatio n of the bacterial [2Fe-2S] ferredoxin is compared with the corresponding metal-bind ing 2Cys2His residues in FD2 of Apd1. There is no corresponding metal-binding site in FD1 of Apd1 (**C**) ^55^Fe radiolabeling of indicated yeast cells expressing HA-Apd1 from a low-copy vector was carried out as in Fig. 3C-E. (**D**) The *apd1Δ* cells from part (C) expressing WT or mutant HA-tagged Apd1 proteins (D1, HA-Apd1-D1; D4, HA-Apd1-D4) from a high-copy vector were subjected to the ^55^Fe radiolabeling-immunoprecipitation assay. (**E-F**) The tagged HA-Apd1-Cap protein was expressed in the indicated (depleted; ↓) yeast ce **l**s. After ^55^Fe radiolabeling, HA-Apd1-Cap (E) and endogenous Leu1 (F) proteins were immunoprecipitated, and ^55^Fe binding was quantified by scintillation counting. Bound ^55^Fe levels were normalized to the respective WT values (5.1 ± 2.3 x 10^3^ and 7.5 ± 4.5 x 10^3^ cpm/g cells, respectively, depicted by dashed horizontal lines). Control experiments involving cells transformed with plasmids lacking Apd1 are depicted as WT– (C,E-F) or ‒Apd1 (D). Values in bar charts represent the mean ± SD, n ≥ 3. For protein expression analysis see Fig. S15B-C.

N-terminally HA-tagged Apd1 expressed from a low-copy plasmid showed effic ie nt complementation in a gallobenzophenone sensitivity assay in *apd1Δ* cells, verifying its functionality (Fig. S13A-B) (40). Upon ^55^Fe radiolabeling and immunoprecipitation with anti-HA beads, HA-Apd1 in WT cells bound a substantial amount of ^55^Fe as estimated by scintillation counting (Fig. 5C). Upon depletion of the early ISC components Nfs1 and Yah1 or of Atm1, ^55^Fe incorporation into HA-Apd1 was substantially diminished, indicating that the bound ^55^Fe was part of a Fe/S cluster. We next depleted cGrxs (using the *cGRX↓aft1Δ* strain) as well as several components of the early (Dre2 and Nbp35) and the late (Nar1, Cia1, and Cia2) CIA system (Fig. 5C, Table S3). Similar to the ISC depletions, deficiency of the cytosolic Fe/S cluster components led to a substantial decrease in ^55^Fe binding in all cases, and was comparative to the loss of ^55^Fe binding to the cytosolic [4Fe-4S] target protein Leu1 (Fig. S12B). A similar ISC dependency and a weaker CIA dependency was observed for HA-Apd1 overexpressed from a high-copy plasmid (Fig. S14). Western blotting indicated lower levels of Apd1 in the ISC mutants during ^55^Fe radiolabeling under low-copy expression (Fig. 5C), yet all levels of Apd1 during high-copy expression were comparable (Fig. S14C).

The presence of a C-terminal tryptophan (Trp) residue in Apd1 may potentially explain its CIA dependence, in contrast to other cytosolic [2Fe-2S] target proteins (39, 40). This residue is responsible for directing Fe/S apoproteins and the adapter protein Lto1 to the CTC (12, 66), and is found in some 20% of known cytosolic-nuclear [4Fe-4S] target apoproteins (39). Indeed, expression of a mutant Apd1 protein lacking either one (HA-Apd1-D1) or four C-terminal residues (HA-Apd1-D4) resulted in approximately 40% or 20%, respectively, of WT ^55^Fe binding (Fig. 5D, bottom). The mutations and the impaired cofactor binding hardly affected Apd1 protein levels (Fig. 5D, top). To further test the influence of the C terminus of Apd1, we masked it by adding a 25-residue long tail (HA-Apd1-Cap) to interfere with Apd1 association with the CTC (Fig. S15C). Like Apd1-D1, HA-Apd1-Cap could not complement Apd1 deficiency in the gallobenzophenone sensitivity assay (Fig. S13B). Despite the lack of functional complementation, HA-Apd1-Cap bound ^55^Fe in a strong ISC-dependent fashion, but in this case ^55^Fe binding in various CIA mutants was at levels observed in wild-type cells, different from Leu1 but similar to canonical [2Fe-2S] proteins (Fig. 5E,F and Fig. S15). The C-terminal tail reduced levels of ^55^Fe binding to Apd1-Cap by nine-fold as compared to the WT protein. Overall, Apd1 showed both a strong ISC and CIA dependency, but upon shielding its C-terminal Trp motif may also bind Fe/S clusters independently of the CIA machinery. The precise molecular mechanisms underlying the special maturation pathway of Apd1 will require dedicated future studies.

### Yeast Bol2 is not involved in [2Fe-2S] cluster maturation of Yap5, Apd1, or CIA proteins

Yeast Bol2 forms a heterodimer with Grx3 or Grx4 via a bridging [2Fe-2S] cluster (20, 21, 23). Bol2-cGrx complexes are proposed to deliver [2Fe-2S] clusters to Aft1-Aft2 (19, 67), although direct *in vivo* evidence is lacking. Bol2 depletion has no major impact on cytosolic [4Fe-4S] protein assembly, and hence is not a CIA component (68). In contrast, studies *in vitro* and in human cells have indicated a role for the human homolog BOLA2 in delivering [2Fe-2S] clusters to the CIA protein CIAPIN1 (25, 28). To clarify the involvement of yeast Bol2 in cytosolic [2Fe-2S] protein maturation apart from its established function in Aft1-Aft2- dependent iron regulation, we determined the ISC and CIA dependence of ^55^Fe binding to Bol2 *in vivo*. Immunoprecipitated HA-tagged Bol2 bound significant amounts of ^55^Fe, which was enhanced three-fold by overexpression of its partner protein Grx4 or, vice versa, was nearly completely abolished by cGrx depletion (Fig. 6A). This result is consistent with biochemical data (28, 69) that only Bol2-cGrx complexes but not Bol2 alone could bind ^55^Fe. Bound ^55^Fe was part of an Fe/S cluster, because it was strongly diminished by depletion of ISC proteins or Atm1 (Fig. 6B). Notably, CIA protein depletion did not affect ^55^Fe binding to Bol2. This result further supports the view that cytosolic [2Fe-2S] protein assembly is generally CIA- independent.

**Figure 6.**
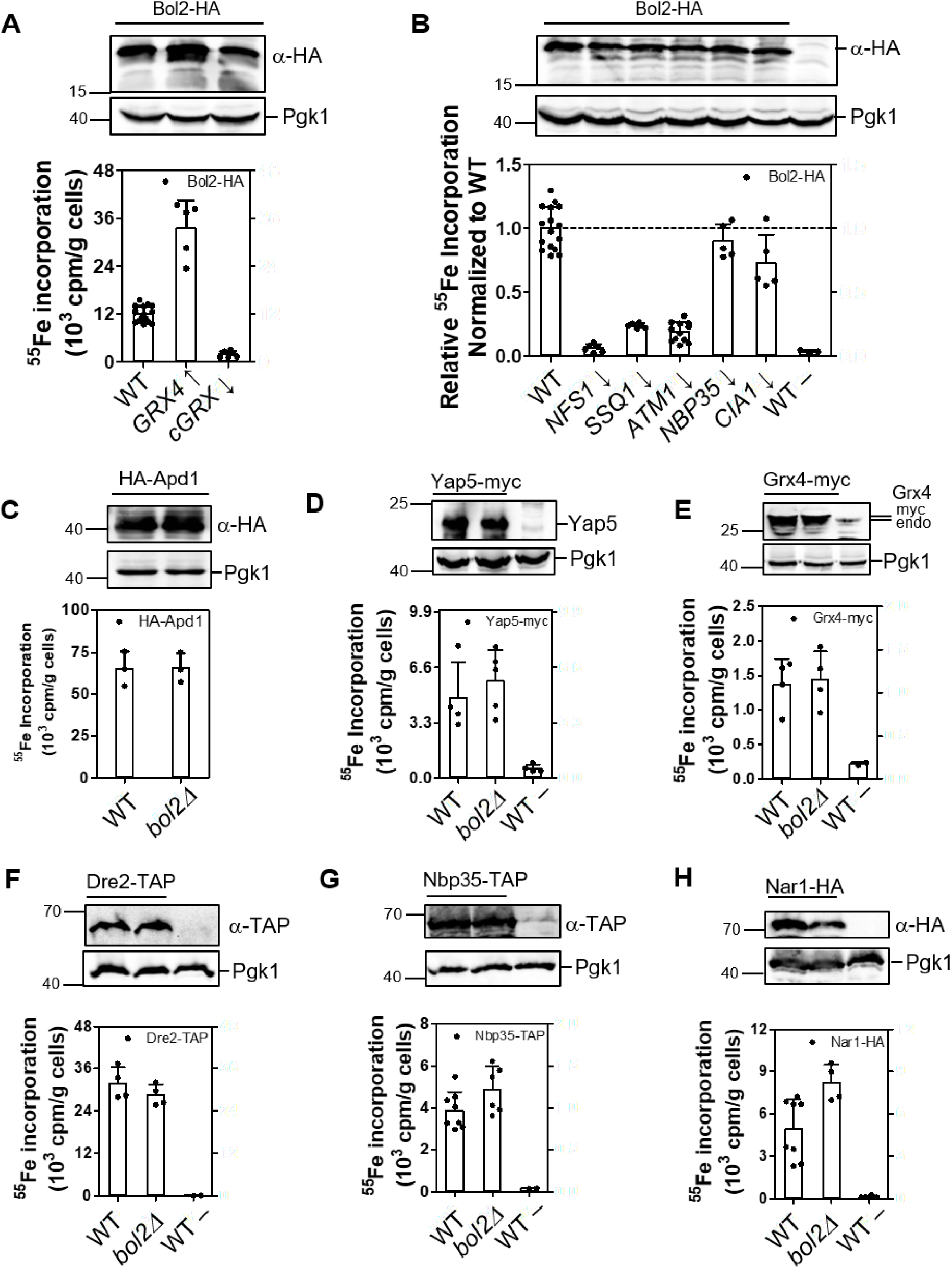
Bol2 is not required for cytosolic [2Fe-2S] protein biogenesis. (**A, B**) Bol2-HA was expressed in wild-type (WT) cells, in the galactose-induced Gal-*GRX4grx3Δ* strain (GRX4↑), and in various glucose-depleted (↓) Gal-strains. ^55^Fe binding of Bol2-HA was quantified as in Fig. 3C-E. (**C-H**) The Bol2 dependence of ^55^Fe binding to indicated cytosolic Fe/S reporter proteins was tested in WT and *bol2Δ* cells followed by the ^55^Fe radiolabeli ng-immunoprecipitation experiment (cf. Fig. 3C-E). The indicated tagged or control (Pgk1) proteins were visualized by immunostaining. WT cells transformed with empty plasmids are depicted as WT–. Values in bar charts represent the mean ± SD, n ≥ 3 (except for control WT– samples in E-G, where n=2).

Reciprocal experiments were performed to determine, if Bol2 influences the assembly of other cytosolic [2Fe-2S] proteins. ^55^Fe binding of HA-Apd1, Yap5-myc, and Grx4-myc was not affected by *BOL2* deletion (Fig. 6C-E). WT levels of ^55^Fe binding to HA-Apd1 provides a striking contrast to the CIA mutants (compare Figs. 5C and 6C). Likewise, the absence of Bol2 did not diminish ^55^Fe binding to the CIA proteins Dre2, Nbp35, or Nar1 (Fig. 6F-H). Together, these results clearly demonstrate that yeast Bol2 is not involved in the maturation of cytosolic [2Fe-2S] or [4Fe-4S] proteins. Hence, the role of yeast Bol2 seems to be confined to Aft1-Aft2- dependent iron regulation, which consistently responds to ISC but not CIA protein depletions. This notion is corroborated by the fact that Bol2 is not essential for cell viability, unlike most CIA proteins (62, 63).

### GSH is required for all cytosolic Fe/S proteins and is linked to Atm1-mediated export

Currently, it is unknown whether GSH is needed for cytosolic [2Fe-2S] protein maturation as it is for [4Fe-4S] proteins (15, 16). Moreover, the exact site of GSH requirement in Fe/S protein biogenesis remains to be determined. However, the dispensable function of the GSH-binding cGrxs in cytosolic Fe/S protein biogenesis (see above) indicated that the essential role of GSH must lie in another step of the pathway. We addressed these open questions by measuring the maturation of various cytosolic Fe/S proteins during GSH depletion in the *gsh1Δ* yeast strain lacking the first biosynthetic gene for GSH (15, 16). GSH depletion severely diminished the ^55^Fe incorporation into the cytosolic [2Fe-2S] proteins HA-Apd1 and Yap5-myc showing that GSH is also essential for this type of cytosolic Fe/S proteins (Fig. 7A,B). As a control, the mitochondria-localized [2Fe-2S] protein Ilv3 did not show any dependence on diminished GSH levels (Fig. 7C). Next, we analyzed which of the Fe/S cluster-containing CIA components may be affected by GSH depletion. Radiolabeling showed near background levels of ^55^Fe binding to Dre2, Nar1-HA, and Nbp35-TAP/HA-Cfd1 upon GSH depletion, clearly defining that all these Fe/S cluster-binding CIA components critically require GSH for maturation (Fig. 7D-E). These results suggest that maturation of both cytosolic [2Fe-2S] and [4Fe-4S] proteins essentially requires GSH. Since the mitochondrial ISC machinery is still unaffected under this level of GSH depletion (15, 16) (Fig. 7C), the Atm1 export step appears to be the most sensitive GSH-dependent reaction in cellular Fe/S protein biogenesis.

**Figure 7.**
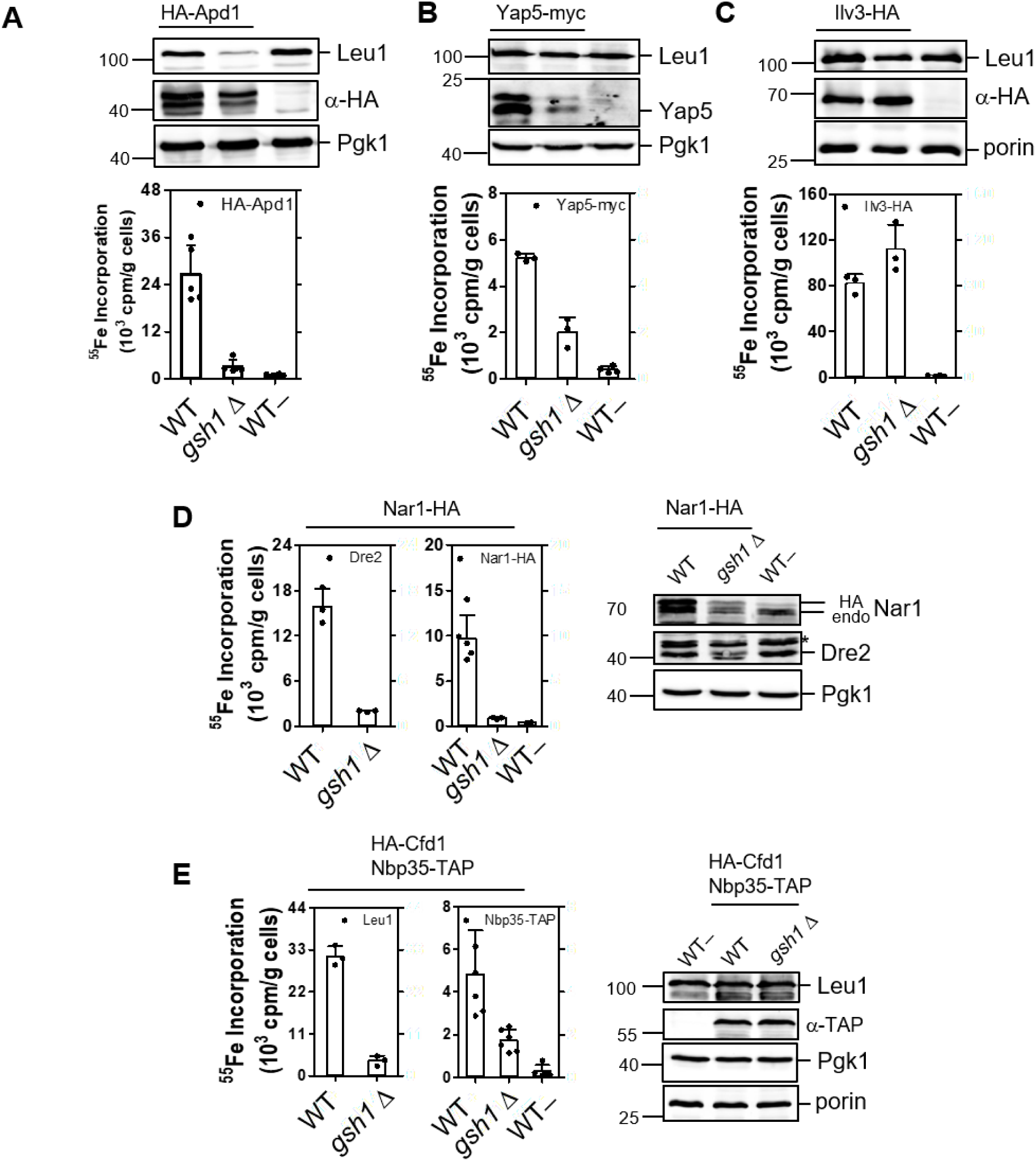
GSH is essential for maturation of cytosolic [2Fe-2S] proteins and Fe/S cluster-binding CIA proteins. Binding of ^55^Fe to indicated [2Fe-2S] target proteins (**A-C**) and CIA components (**D-E**) in wild-type (WT; YPH500) and *gsh1Δ* yeast strains was analyzed as in Fig. 3C-E. WT cells transformed with yeast plasmids lacking epitope tags (WT–) served as controls. The indicated tagged or native proteins and the loading controls (Pgk1 or porin) were visuali zed by immunostaining. Values represent the mean ± SD, n ≥ 3. *, denotes a non-specific band.

## Discussion

Work from nearly two decades has elucidated the eukaryotic CIA pathway with its more than 10 components for the proper synthesis, trafficking, and insertion of [4Fe-4S] clusters into cytosolic, ER-associated, and nuclear [4Fe-4S] target proteins in various eukaryotes (Table S1) (1, 2). A major gap in our knowledge of cellular Fe/S protein assembly is the origin, traffick ing, and apoprotein insertion of cytosolic-nuclear [2Fe-2S] clusters. Only relatively few such proteins are known in the eukaryotic cytosol, yet they perform important functions in, e.g., the early CIA machinery, iron homeostasis, redox regulation, and metabolic processes (see Supplemental Table 2 in (4)). In contrast to mitochondria, the eukaryotic cytosol does not contain a known [2Fe-2S] cluster scaffold protein, and hence it has remained unclear, if [2Fe-2S] clusters need to be *de novo* synthesized in the cytosol. Further, there are conflicting reports in the current literature, which of the mitochondrial (ISC) or particularly cytosolic (CIA) Fe/S protein assembly system are needed to mature these [2Fe-2S] proteins (see Introduction).

By analyzing most of the cytosolic [2Fe-2S] cluster-binding proteins in both yeast and human cells by *in vivo* ^55^Fe radiolabeling-immunoprecipitation experiments, we report that, with one exception, all tested cytosolic [2Fe-2S] proteins are matured independently of the known CIA components and cGrxs, yet strictly depend on the early-acting ISC components and the Atm1-ABCB7-facilitated X-S export from mitochondria (Fig. 8, blue box). Thus, cytosolic [2Fe-2S] cluster maturation characteristically differs from that of cytosolic [4Fe-4S] proteins which require both mitochondrial and cytosolic biogenesis systems (70) (Fig. 8, green box). Apparently, in lieu of an obvious cytosolic [2Fe-2S] scaffold, this cluster type or its precursor is synthesized in and provided by mitochondria, and then can be inserted into apoproteins without the essential assistance of the known CIA proteins and cGrxs. Our findings do not exclude, however, the existence of so far unknown factors facilitating the cytosolic assembly process. The predominant CIA independence of cytosolic [2Fe-2S] protein biogenesis reported here fits well to the CIA independence of iron regulation in fungi essentially involving the cGrxs (18–22, 33, 63). In marked contrast, iron regulation in mammals depends on, e.g., the [4Fe-4S] protein IRP1, and thus is CIA-responsive (11).

**Figure 8.**
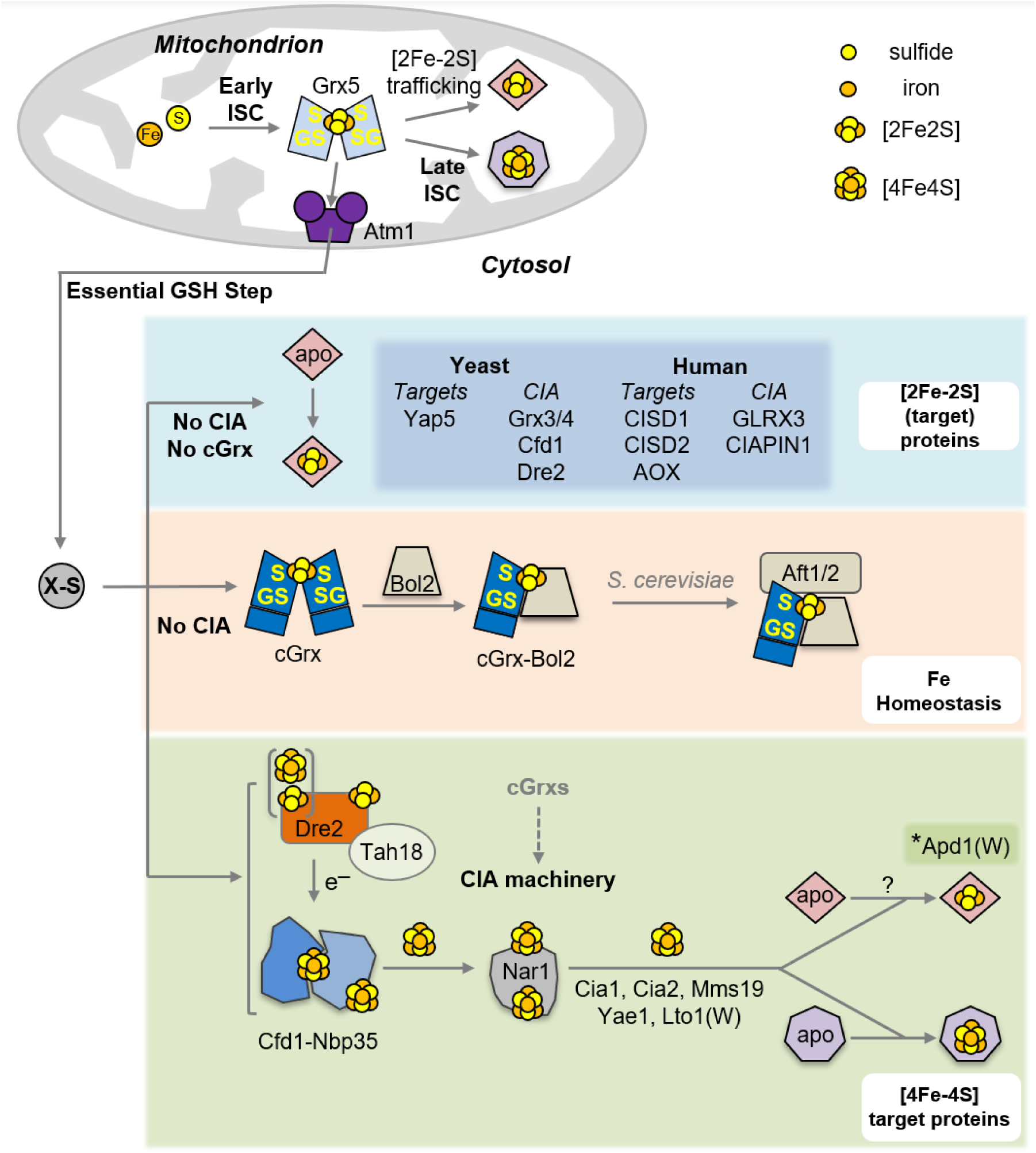
Working model for [2Fe-2S] and [4Fe-4S] protein biogenesis in the eukaryotic cytosol. In mitochondria, the early and late ISC pathways, connected by the monothiol glutaredoxin Grx5, are essential for the maturation of organellar [2Fe-2S] and [4Fe-4S] proteins (top). The early ISC system and Grx5 are also required for generating the sulfur- and possibly iron-containing compound X-S for its export by the ABC transporter Atm1 thereby assisting the CIA machinery in cytosolic [4Fe-4S] target protein assembly (green box). In this work, the so far unclear maturation pathways for cytosolic [2Fe-2S] proteins, as well as the roles of glutathione (GSH) and the cytosolic monothiol glutaredoxins (cGrxs) in general cytosolic Fe/S protein maturation were elucidated *in vivo* in yeast and human cells. Our studies, together with published work, suggest that cytosolic [2Fe-2S] target proteins and the three [2Fe-2S] cluster-binding CIA proteins (see dark blue box for tested examples) require the early ISC machinery, Atm1, and GSH for maturation, yet are independent of any of the known CIA components and the cGrxs (light blue box). *, One striking exception is yeast Apd1, which depends on the CIA machinery for [2Fe-2S] cluster assembly (dotted arrow), due to its CIA targeting complex CTC) binding via its C-terminal tryptophan (W). The ISC dependence and CIA independence also applies to [2Fe-2S] cluster maturation of cGrxs and cGrx-Bol2 complexes (orange box). In yeast, the cGrx-Bol2 holo-complex performs a crucial, CIA-independent role in cellular Fe regulation by [2Fe-2S] cluster-dependent attenuation of the Aft1-Aft2 transcription factors. Yeast and human cGrxs play a dispensable, mechanistically still unclear role in [4Fe-4S] protein maturation by the CIA pathway (dashed arrow in green box). cGrxs genetically and physically interact with yeast Dre2 and human CIAPIN1, yet this property is not essential for CIA function and cell viability. The insertion of Fe/S clusters into the CIA components Cfd1, Cfd1-Nbp35, and yeast Dre2 (with one [2Fe-2S] and one [4Fe-4S] cluster) or human CIAPIN1 (two [2Fe-2S] clusters) is dependent on X-S but does not require cGrxs. GS, Grx-bound, cluster-coordinating glutathione.

A curious exception to the CIA-independent pathway of cytosolic [2Fe-2S] proteins is yeast Apd1, which shows a strong CIA dependency. A C-terminal tryptophan, often seen in [4Fe-4S] target proteins or adapters interacting with the CTC like Lto1, is likely responsible for guiding Apd1 to the CTC for proper Fe/S cluster insertion (Fig. 8, dotted arrow) (12, 39). Interestingly, Apd1 with an extended C-terminus (Apd1-Cap) could bind Fe/S clusters in a CIA-independent fashion, yet was not functionally active. As all of the CIA components have previously been described to support [4Fe-4S] protein maturation (1, 2), Apd1’s maturation pathway appears to be more complex than that of the other [2Fe-2S] proteins which receive their cofactor in a CIA- independent way. This might equally apply to the Rieske-type cluster containing NEVERLAND and choline monooxygenase proteins, which like Apd1 have a conserved C-terminal tryptophan (39). Hence, further studies are required to discriminate whether the CIA system can also directly assemble a [2Fe-2S] cluster, or whether alternatively Apd1 first receives a [4Fe-4S] cluster at the CIA targeting complex, that is subsequently rearranged into a [2Fe-2S] cluster.

Our study furthermore suggests that the long-known essential requirement of GSH for extra-mitochondrial Fe/S protein biogenesis is primarily connected to the Atm1-ABCB7 export reaction rather than to the GSH-dependent function of cGrxs (Fig. 8) (15, 16). Currently, the chemical composition of X-S and therefore its role in the CIA pathway remains elusive. The findings from our current *in vivo* studies showing no obvious requirement of cytosolic factors for most [2Fe-2S] protein maturation and from an *in vitro* reconstitution of the export pathway (71) would be consistent with the proposed export of GSH-containing [2Fe-2S] clusters from mitochondria by Atm1-ABCB7 (9). Such a mechanism would provide [2Fe-2S] clusters for cytosolic apoproteins, including early CIA components, without the need for cytosolic *de novo* synthesis as suggested by other work in the field (Fig. 8) (72, 73). Clearly, further insight into this aspect of cytosolic Fe/S protein biogenesis depends on the molecular identification of X-S *in vivo*. In mammals, NEET proteins have been claimed to be involved in a hand-off of [2Fe-2S] clusters from the mitochondrial inter membrane space (CISD3) to the cytosol (CISD1) via porin (VDAC1) channels (74). It remains puzzling why the NEETs are not found in, e.g., fungi, and why only CISD3 and not CISD3 is essential (75).

Similar to our findings, the loading of Fe/S clusters into a purified [2Fe-2S] target apoprotein (a mitochondrial ferredoxin) was observed to be dependent on mitochondria exporting an iron- and sulfur-containing species (41). However, contrary to our *in vivo* findings, a dependence on the CIA components Dre2 and Cfd1 was reported for this *in vitro* approach, similar as for *in vitro* [4Fe-4S] protein maturation (71). In our *in vivo* radiolabeling study, depletion of Dre2 or CIAPIN1 was found to maintain or even increase ^55^Fe binding to most [2Fe-2S] target proteins, clearly indicating a CIA-independent flow of [2Fe-2S] clusters in the cytosol *in vivo*. This discrepancy needs to be resolved in future work.

Although the cytosolic monothiol glutaredoxins have been shown *in vitro* to transfer [2Fe-2S] clusters to proteins like Dre2 (25, 27, 28) and [2Fe-2S] target proteins, our experiments in both yeast and human cells showed that such a transfer function is dispensable or can be completely bypassed. Our observations are consistent with the fact that cGrxs are not essential in various eukaryotes. Reported essentiality in some species seems to be connected to their additional role in iron regulation as shown here for *S. cerevisiae* cGrxs (Table S2; see below). Interestingly, the cGrx functional bypass occurs despite cGrx and Dre2 homologs interacting in various organisms (25, 33). Human GLRX3 has also been found to interact with components of the late CIA machinery (CIAO1 and CIAO2B) (11, 25), and may be connected to the [4Fe-4S] cluster defects in *GLRX3* KO cells. Despite the potentially multiple influences of cGrxs on the CIA system, either to increase Fe/S cluster trafficking efficiency or to fulfil a so far unknown function, it is clear from our work that their presence is not essential for a critical flow of [4Fe-4S] clusters via the CIA system to cytosolic and nuclear target proteins. A similar functional bypass has also been reported for the monothiol Grx5 of the mitochondrial ISC system in yeast, despite its proposed role in connecting early and late ISC components (Fig. 8) (76).

The essential requirement of cGrxs for cell viability of some single-celled eukaryotes like *S. cerevisiae* (W303) (18) and *Aspergillus fumigatus* (33) appears to be intimately connected to the function of these proteins as sensors of iron homeostasis (Figs. 5, 8). Concomitant gene deletion of both cGrxs and the corresponding transcriptional regulators of iron uptake, i.e. *S. cerevisiae* Aft1 or *A. fumigatus* SreA, suppressed the lethality of cGrx gene deletion (this work; (33)). Along these lines, the critical function of cGrxs in iron homeostasis can be bypassed by other genetic suppressors, which rewire cellular networks such as pH sensing and nutritional signaling (56). Permanently deactivating these sensors leads to iron overload, which is lethal especially in the presence of oxygen by formation of damaging reactive oxygen species (56). In mammals, where GLRX3 has no known direct role in iron regulation, its gene deletion is lethal during mouse embryonic development (77, 78), but the molecular explanation remains to be worked out. Likewise, it needs to be determined why this essentiality does not translate to tissue-specific gene knockouts, e.g., in mammary-, heart-, or kidney-derived cell types or to our CRISPR-mediated gene knockout (Table S2). Finally, further work is warranted to fully disentangle the iron homeostasis and [4Fe-4S] protein biogenesis defects in cells lacking cGrxs.

## Materials and Methods

### Yeast strains and plasmids

*Saccharomyces cerevisiae* strains (W303-1A and YPH500 backgrounds) used are listed in Table S3. Yeast cells were grown in either rich (YP) or minimal (SC) medium with all required supplements and 2% glucose (D) or galactose (Gal) (79). Plasmids used for the expression of proteins in yeast are listed in Table S4A.

### Human tissue culture and methods

HeLa and HEK293 cell lines were maintained and manipulated as described in the Supplemental Methods. Transfection of siRNAs (Table S5A) and plasmids (Table S4B-C) into human cells by electroporation or chemical methods was carried out by standard techniques as described also in the Supplemental Methods (48). Specific primers used for cloning are listed in Tables S5B-C. CRISPR Cas9-mediated gene knock-out or knock-in was performed according to published protocols using guide RNAs (gRNA) described in Table S5C (53).

### Miscellaneous Methods

Home-made and commercial antibodies are listed in Table S6. Manipulation of DNA and PCR (80), immunological techniques (81), and protein determination by the BCA method (Thermo Scientific) were carried out according to standard procedures. Transformation of yeast cells (82) and *in vivo* labeling of iron-binding proteins with ^55^FeCl3 (Perkin Elmer) and immunoprecipitation of ^55^Fe/S proteins and scintillation counting were performed as published for yeast (59) and human cells (47, 48). The expression of *GFP* from the *FET3* promoter was performed as published (58). Gallobenzophenone growth sensitivity of *apd1Δ* cells was performed as described (40). New DNA constructs were confirmed by Sanger sequencing.

## Acknowledgements

We thank Marlina Indrayani, Manuel Müller, and Brigitte Niggeme yer for their technical assistance. This work was supported by a Marie Skłodowska-Curie Fellowship 659325 (to J. J. B.), grants from Deutsche Forschungsgemeinschaft (DFG) within SPP 1927 (to J. J. B. and A. J. P.), and financial support (to R. L.) from the DFG (Koselleck grant, LI 415/5, SFB 987 (R. L. and U. M.), SPP 1710, and SPP 1927).

## The abbreviations used are

Fe/S: iron-sulfur
ISC: iron-sulfur cluster assembly
CIA: cytosolic iron-sulfur protein assembly
Grx: glutaredoxin
cGrx: cytosolic monothio l glutaredoxin
GSH: glutathione
GSSSG: oxidized glutathione persulfide
Moco: molybdenum cofactor
CTC: CIA targeting complex
FD: ferredoxin-like domain
Aox: aldehyde oxidase
CTD: C-terminal domain
KO: knock out
gRNA: guide RNA
siRNA: small interfering RNA
shRNA: small hairpin RNA
TEV: tobacco edge virus
EGFP: enhanced green fluoresce nt protein.

## Author Contributions

J.J.B., O.S., U.M., A.J.P., and R.L. designed research; J.J.B., O.S., M.S., R.R., F.S., C.M.B., and U.M. performed research; J.J.B., O.S., A.J.P., U.M. and R.L. analyzed data; all authors read and reviewed the paper; J.J.B, O.S., and R.L. wrote the paper.

## References

1. Braymer JJ, Freibert SA, Rakwalska-Bange M, & Lill R (2021) Mechanistic concepts of iron-sulfur protein biogenesis in Biology. Biochim Biophys Acta Mol Cell Res 1868(1):118863.

2. Ciofi-Baffoni S, Nasta V, & Banci L (2018) Protein networks in the maturation of human iron-sulfur proteins. Metallomics 10(1):49–72.

3. Maio N & Rouault TA (2020) Outlining the Complex Pathway of Mammalian Fe-S Cluster Biogenesis. Trends Biochem Sci 45(5):411–426.

4. Lill R & Freibert SA (2020) Mechanisms of Mitochondrial Iron-Sulfur Protein Biogenesis. Annu Rev Biochem 89:471–499.

5. Dutkiewicz R & Nowak M (2018) Molecular chaperones involved in mitochondrial iron-sulfur protein biogenesis. J Biol Inorg Chem 23(4):569–579.

6. Weiler BD, et al. (2020) Mitochondrial [4Fe-4S] protein assembly involves reductive [2Fe-2S] cluster fusion on ISCA1-ISCA2 by electron flow from ferredoxin FDX2. Proc Natl Acad Sci U S A 117(34):20555–20565.

7. Netz DJ, Mascarenhas J, Stehling O, Pierik AJ, & Lill R (2014) Maturation of cytosolic and nuclear iron-sulfur proteins. Trends Cell Biol 24(5):303–312.

8. Srinivasan V, Pierik AJ, & Lill R (2014) Crystal structures of nucleotide-free and glutathione-bound mitochondrial ABC transporter Atm1. Science 343(6175):1137–1140.

9. Li P, et al. (2022) Structures of Atm1 provide insight into [2Fe-2S] cluster export from mitochondria. Nat Commun 13(1):4339.

10. Bargagna B, Matteucci S, Ciofi-Baffoni S, Camponeschi F, & Banci L (2023) Unraveling the mechanism of [4Fe-4S] cluster assembly on the N-terminal cluster binding site of NUBP1. Protein Sci 32(5):e4625.

11. Stehling O, et al. (2013) Human CIA2A-FAM96A and CIA2B-FAM96B integrate iron homeostasis and maturation of different subsets of cytosolic-nuclear iron-sulfur proteins. Cell Metab 18(2):187–198.

12. Paul VD, et al. (2015) The deca-GX_3_ proteins Yae1-Lto1 function as adaptors recruiting the ABC protein Rli1 for iron-sulfur cluster insertion. eLife 4:e08231.

13. Kassube SA & Thoma NH (2020) Structural insights into Fe-S protein biogenesis by the CIA targeting complex. Nat Struct Mol Biol.

14. Maione V, Grifagni D, Torricella F, Cantini F, & Banci L (2020) CIAO3 protein forms a stable ternary complex with two key players of the human cytosolic iron-sulfur cluster assembly machinery. J Biol Inorg Chem 25(3):501–508.

15. Sipos K, et al. (2002) Maturation of cytosolic iron-sulfur proteins requires glutathione. J. Biol. Chem. 277:26944–26949.

16. Kumar C, et al. (2011) Glutathione revisited: a vital function in iron metabolism and ancillary role in thiol-redox control. Embo J 30(10):2044–2056.

17. Schaedler TA, et al. (2014) A Conserved Mitochondrial ATP-Binding Cassette Transporter Exports Glutathione Polysulfide for Cytosolic Metal Cofactor Assembly. J Biol Chem 289(34):23264–23274.

18. Mühlenhoff U, et al. (2010) Cytosolic monothiol glutaredoxins function in intracellular iron sensing and trafficking via their bound iron-sulfur cluster. Cell Metab 12(4):373–385.

19. Poor CB, et al. (2014) Molecular mechanism and structure of the Saccharomyces cerevisiae iron regulator Aft2. Proc Natl Acad Sci U S A 111(11):4043–4048.

20. Muhlenhoff U, et al. (2020) Glutaredoxins and iron-sulfur protein biogenesis at the interface of redox biology and iron metabolism. Biol Chem 401(12):1407–1428.

21. Talib EA & Outten CE (2021) Iron-sulfur cluster biogenesis, trafficking, and signaling: Rolesfor CGFS glutaredoxins and BolA proteins. Biochim Biophys Acta Mol Cell Res 1868(1):118847.

22. Kumanovics A, et al. (2008) Identification of FRA1 and FRA2 as genes involved in regulating the yeast iron regulon in response to decreased mitochondrial iron-sulfur cluster synthesis. J Biol Chem 283:10276–10286.

23. Hati D, et al. (2023) Iron homeostasis proteins Grx4 and Fra2 control activity of the Schizosaccharomyces pombe iron repressor Fep1 by facilitating [2Fe-2S] cluster removal. J Biol Chem 299(12):105419.

24. Haunhorst P, et al. (2013) Crucial function of vertebrate glutaredoxin 3 (PICOT) in iron homeostasis and hemoglobin maturation. Mol Biol Cell 24(12):1895–1903.

25. Frey AG, Palenchar DJ, Wildemann JD, & Philpott CC (2016) A Glutaredoxin.BolA Complex Serves as an Iron-Sulfur Cluster Chaperone for the Cytosolic Cluster Assembly Machinery. J Biol Chem 291(43):22344–22356.

26. Camponeschi F, Prusty NR, Heider SAE, Ciofi-Baffoni S, & Banci L (2020) GLRX3 Acts as a [2Fe-2S] Cluster Chaperone in the Cytosolic Iron-Sulfur Assembly Machinery Transferring [2Fe-2S] Clusters to NUBP1. J Am Chem Soc 142(24):10794–10805.

27. Banci L, et al. (2015) N-terminal domains mediate [2Fe-2S] cluster transfer from glutaredoxin-3 to anamorsin. Nat Chem Biol 11(10):772–778.

28. Banci L, Camponeschi F, Ciofi-Baffoni S, & Muzzioli R (2015) Elucidating the Molecular Function of Human BOLA2 in GRX3-Dependent Anamorsin Maturation Pathway. J Am Chem Soc 137(51):16133–16143.

29. Rietzschel N, Pierik AJ, Bill E, Lill R, & Muhlenhoff U (2015) The basic leucine zipper stress response regulator Yap5 senses high-iron conditions by coordination of [2Fe-2S] clusters. Mol Cell Biol 35(2):370–378.

30. Linder T (2014) CMO1 encodes a putative choline monooxygenase and is required for the utilization of choline as the sole nitrogen source in the yeast Scheffersomyces stipitis (syn. Pichia stipitis). Microbiology (Reading) 160(Pt 5):929–940.

31. Rosenbach H, et al. (2021) The Asp1 pyrophosphatase from S. pombe hosts a [2Fe-2S](2+) cluster in vivo. J Biol Inorg Chem 26(1):93–108.

32. Kim HJ, Lee KL, Kim KD, & Roe JH (2016) The iron uptake repressor Fep1 in the fission yeast binds Fe-S cluster through conserved cysteines. Biochem Biophys Res Commun 478(1):187–192.

33. Misslinger M, et al. (2019) The monothiol glutaredoxin GrxD is essential for sensing iron starvation in Aspergillus fumigatus. PLoS Genet 15(9):e1008379.

34. Netz DJ, et al. (2010) Tah18 transfers electrons to Dre2 in cytosolic iron-sulfur protein biogenesis. Nat Chem Biol 6(10):758–765.

35. Zhang Y, Yang C, Dancis A, & Nakamaru-Ogiso E (2017) EPR studies of wild type and mutant Dre2 identify essential [2Fe--2S] and [4Fe--4S] clusters and their cysteine ligands. J Biochem 161(1):67–78.

36. Netz DJ, et al. (2016) The conserved protein Dre2 uses essential [2Fe-2S] and [4Fe-4S] clusters for its function in cytosolic iron-sulfur protein assembly. Biochem J 473(14):2073–2085.

37. Matteucci S, et al. (2021) In-cellulo Mössbauer and EPR studies bring new evidences to the long-standing debate on the iron-sulfur cluster binding in human anamorsin. Angew Chem Int Ed Engl.

38. Ferecatu I, et al. (2014) The diabetes drug target MitoNEET governs a novel trafficking pathway to rebuild an Fe-S cluster into cytosolic aconitase/iron regulatory protein 1. J Biol Chem 289(41):28070–28086.

39. Marquez MD, et al. (2023) Cytosolic iron-sulfur protein assembly systemidentifies clients by a C-terminal tripeptide. Proc Natl Acad Sci U S A 120(44):e2311057120.

40. Stegmaier K, et al. (2019) Apd1 and Aim32 Are Prototypes of Bishistidinyl-Coordinated Non-Rieske [2Fe-2S] Proteins. J Am Chem Soc 141(14):5753–5765.

41. Pandey AK, Pain J, Dancis A, & Pain D (2019) Mitochondria export iron-sulfur and sulfur intermediates to the cytoplasm for iron-sulfur cluster assembly and tRNA thiolation in yeast. J Biol Chem 294(24):9489–9502.

42. Song D & Lee FS (2008) A role for IOP1 in mammalian cytosolic iron-sulfur protein biogenesis. J. Biol. Chem. 283(14):9231–9238.

43. Song D & Lee FS (2011) Mouse knock-out of IOP1 protein reveals its essential role in mammalian cytosolic iron-sulfur protein biogenesis. J Biol Chem 286(18):15797–15805.

44. Bastow EL, Bych K, Crack JC, Le Brun NE, & Balk J (2017) NBP35 interacts with DRE2 in the maturation of cytosolic iron-sulphur proteins in Arabidopsis thaliana. Plant J 89(3):590–600.

45. Terao M, Garattini E, Romao MJ, & Leimkuhler S (2020) Evolution, expression, and substrate specificities of aldehyde oxidase enzymes in eukaryotes. J Biol Chem 295(16):5377–5389.

46. Schumann S, et al. (2008) The mechanism of assembly and cofactor insertion into Rhodobacter capsulatus xanthine dehydrogenase. J Biol Chem 283(24):16602–16611.

47. Stehling O, Paul VD, Bergmann J, Basu S, & Lill R (2018) Biochemical Analyses of Human Iron–Sulfur Protein Biogenesis and of Related Diseases. Methods Enzymol. 599:227–263.

48. Stehling O, et al. (2018) Function and crystal structure of the dimeric P-loop ATPase CFD1 coordinating an exposed [4Fe-4S] cluster for transfer to apoproteins. Proc Natl Acad Sci U S A 115(39):E9085–E9094.

49. Karmi O, et al. (2018) The unique fold and lability of the [2Fe-2S] clusters of NEET proteins mediate their key functions in health and disease. J Biol Inorg Chem 23(4):599–612.

50. Stehling O, et al. (2012) MMS19 Assembles Iron-Sulfur Proteins Required for DNA Metabolism and Genomic Integrity. Science 337(6091):195–199.

51. Soler N, et al. (2012) A S-adenosylmethionine methyltransferase-like domain within the essential, Fe-S-containing yeast protein Dre2. FEBS J 279(12):2108–2119.

52. Banci L, et al. (2013) Human anamorsin binds [2Fe-2S] clusters with unique electronic properties. J Biol Inorg Chem 18(8):883–893.

53. Ran FA, et al. (2013) Genome engineering using the CRISPR-Cas9 system. Nat Protoc 8(11):2281–2308.

54. Paul VD & Lill R (2015) Biogenesis of cytosolic and nuclear iron-sulfur proteins and their role in genome stability. Biochim Biophys Acta 1853(6):1528–1539.

55. Pujol-Carrion N, Belli G, Herrero E, Nogues A, & de la Torre-Ruiz MA (2006) Glutaredoxins Grx3 and Grx4 regulate nuclear localisation of Aft1 and the oxidative stress response in Saccharomyces cerevisiae. J Cell Sci 119(Pt 21):4554–4564.

56. Li G, Nanjaraj Urs AN, Dancis A, & Zhang Y (2022) Genetic suppressors of Deltagrx3 Deltagrx4, lacking redundant multidomain monothiol yeast glutaredoxins, rescue growth and iron homeostasis. Biosci Rep 42(6).

57. Janke C, et al. (2004) A versatile toolbox for PCR-based tagging of yeast genes: new fluorescent proteins, more markers and promoter substitution cassettes. Yeast 21(11):947–962.

58. Molik S, Lill R, & Mühlenhoff U (2007) Methods for studying iron metabolism in yeast mitochondria. Methods Cell. Biol. 80:261–280.

59. Pierik AJ, Netz DJA, & Lill R (2009) Analysis of iron-sulfur protein maturation in eukaryotes. . Nat. Protoc. 4:753–766.

60. Yamaguchi-Iwai Y, Dancis A, & Klausner RD (1995) AFT1: a mediator of iron regulated transcriptional control in Saccharomyces cerevisiae. Embo J 14(6):1231–1239.

61. Chen OS, et al. (2004) Transcription of the yeast iron regulon responds not directly to iron but rather to iron-sulfur cluster biosynthesis. J Biol Chem 279:29513–29518.

62. Rutherford JC, et al. (2005) Activation of the iron regulon by the yeast Aft1/Aft2 transcription factors depends on mitochondrial but not cytosolic iron-sulfur protein biogenesis. J Biol Chem 280(11):10135–10140.

63. Hausmann A, Samans B, Lill R, & Muhlenhoff U(2008) Cellular and Mitochondrial Remodeling upon Defects in Iron-Sulfur Protein Biogenesis. J Biol Chem 283(13):8318–8330.

64. Tang HM, et al. (2015) Loss of APD1 in yeast confers hydroxyurea sensitivity suppressed by Yap1p transcription factor. Sci Rep 5:7897.

65. Varadi M, et al. (2022) AlphaFold Protein Structure Database: massively expanding the structural coverage of protein-sequence space with high-accuracy models. Nucleic Acids Res 50(D1):D439–D444.

66. Upadhyay AS, et al. (2014) Viperin is an iron-sulfur protein that inhibits genome synthesis of tick-borne encephalitis virus via radical SAM domain activity. Cell Microbiol 16(6):834–848.

67. Li H & Outten CE (2019) The conserved CDC motif in the yeast iron regulator Aft2 mediates iron-sulfur cluster exchange and protein-protein interactions with Grx3 and Bol2. J Biol Inorg Chem 24(6):809–815.

68. Uzarska MA, et al. (2016) Mitochondrial Bol1 and Bol3 function as assembly factors for specific iron-sulfur proteins. Elife 5:e16673.

69. Li H, Mapolelo DT, Randeniya S, Johnson MK, & Outten CE (2012) Human glutaredoxin 3 forms [2Fe-2S]-bridged complexes with human BolA2. Biochemistry 51(8):1687–1696.

70. Kispal G, Csere P, Prohl C, & Lill R (1999) The mitochondrial proteins Atm1p and Nfs1p are essential for biogenesis of cytosolic Fe/S proteins. Embo J 18(14):3981–3989.

71. Pandey AK, Pain J, J B, Dancis A, & Pain D (2023) Essential mitochondrial role in iron-sulfur cluster assembly of the cytoplasmic isopropylmalate isomerase Leu1 in Saccharomyces cerevisiae. Mitochondrion 69:104–115.

72. Kim KS, Maio N, Singh A, & Rouault TA (2018) Cytosolic HSC20 integrates de novo iron-sulfur cluster biogenesis with the CIAO1-mediated transfer to recipients. Hum Mol Genet 27(5):837–852.

73. Patel SJ, et al. (2019) A PCBP1-BolA2 chaperone complex delivers iron for cytosolic [2Fe-2S] cluster assembly. Nat Chem Biol 15(9):872–881.

74. Karmi O, et al. (2022) A VDAC1-mediated NEET protein chain transfers [2Fe-2S] clusters between the mitochondria and the cytosol and impacts mitochondrial dynamics. Proc Natl Acad Sci U S A 119(7).

75. Sengupta S, et al. (2018) Phylogenetic analysisof the CDGSH iron-sulfur binding domain reveals its ancient origin. Sci Rep 8(1):4840.

76. Uzarska MA, Dutkiewicz R, Freibert SA, Lill R, & Muhlenhoff U(2013) The mitochondrial Hsp70 chaperone Ssq1 facilitates Fe/S cluster transfer from Isu1 to Grx5 by complex formation. Mol Biol Cell 24(12):1830–1841.

77. Cha H, et al. (2008) PICOT is a critical regulator of cardiac hypertrophy and cardiomyocyte contractility. J Mol Cell Cardiol 45(6):796–803.

78. Cheng NH, et al. (2011) A mammalian monothiol glutaredoxin, Grx3, is critical for cell cycle progression during embryogenesis. FEBS J 278(14):2525–2539.

79. Sherman F (2002) Getting started with yeast. Methods Enzymol. 350:3–41.

80. Sambrook J & Russell DW (2001) Molecular cloning: A laboratory manual (Cold Spring Harbor Press, Cold Spring Harbor, NY) 3rd edition Ed.

81. Harlow E & Lane D (1998) Using Antibodies: A Laboratory Manual (Cold Spring Harbor Laboratory, Cold Spring Harbor, NY).

82. Gietz RD & Woods RA (2002) Transformation of yeast by lithium acetate/single-stranded carrier DNA/polyethylene glycol method. Methods Enzymol 350:87–96.

