## Supplemental for "Requirements for the Biogenesis of [2Fe-2S] Proteins in the Human and Yeast Cytosol"

**Running Title**: Cytosolic [2Fe-2S] cluster trafficking *in vivo*

Joseph J. Braymer[a,b], Oliver Stehling[a,b], Martin Stümpfig[a,b], Ralf Rösser[a,b], Farah Spantgar[a,b], Catharina M. Blinn[c], Ulrich Mühlenhoff[a,b], Antonio J. Pierik[c], Roland Lill[a,b,#]

a. Institut für Zytobiologie und Zytopathologie, Fachbereich Medizin, Philipps-Universität Marburg, Karl-von-Frisch-Straße 14, 35032 Marburg, Germany
b. Zentrum für Synthetische Mikrobiologie Synmikro, Philipps-Universität Marburg, Karl-von-Frisch-Straße 14, 35032 Marburg Germany
c. Department of Chemistry, RPTU Kaiserslautern-Landau, 67663 Kaiserslautern, Germany

[#] To whom correspondence should be addressed:

**Contents**:

**Supplemental Figures**

Figure S1: General overview of eukaryotic Fe/S cluster biogenesis

Figure S2: Supporting data for $^{55}$Fe incorporation into Aox

Figure S3: Supporting data for $^{55}$Fe incorporation into CISD2

Figure S4: Supporting data for $^{55}$Fe incorporation into ABCE1

Figure S5: Supporting data for $^{55}$Fe incorporation into endogenous CIAPIN1 in RNAi experiments

Figure S6: Supporting data for $^{55}$Fe incorporation into endogenous CIAPIN1 in *GLRX3* KO cells

Figure S7: Supporting data for $^{55}$Fe incorporation into over-expressed CIAPIN1 in KO cells

Figure S8: Supporting data for $^{55}$Fe incorporation into Aox, CISD1/2, ABCE1 in *GLRX3* KO cells

Figure S9: Dre2 independence of the CIA system

Figure S10: Full Western blot for Figure 4B

Figure S11: 55Fe incorporation into Yap5-myc

Figure S12: Supporting data for low-copy p416-HA-APD1 expression in ISC and CIA mutant strains

Figure S13: Western blotting and complementation experiments with various Apd1 constructs

Figure S14: Supporting data for high-copy p426-HA-APD1 expression in ISC and CIA mutant strains

Figure S15: Supporting data for low-copy p416-HA-APD1-Cap expression in ISC and CIA mutant strains

**Supplemental Methods**

**Supplemental Tables**

Table S1: Nomenclature of yeast and human proteins involved in Fe/S protein assembly

Table S2: Deletion phenotype of cytosolic monothiol glutaredoxins (cGrxs) studied to date

Table S3: Yeast strains used in this study

Table S4A-B: Plasmid constructs used in this study

Table S4C: CRISPR-related plasmid constructs used for analyses in mammalian cells

Table S4D: shRNA plasmid constructs used for gene silencing in mammalian cells

Table S5A: RNAi sequences

Table S5B: PCR primers used for site-directed mutagenesis of human ABCB7

Table S5C: Guide RNA sequence for CRISPR-Cas9

Table S6: Antibodies used in this study

**Supplemental Figures**

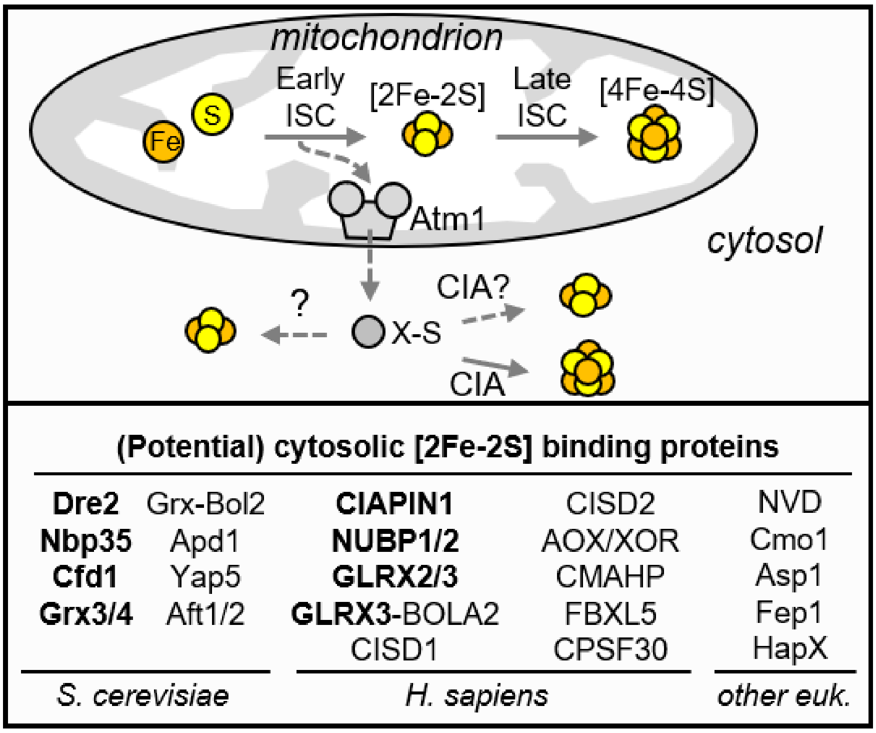

**Figure S1:** Simplified model for maturation of cytosolic-nuclear Fe/S proteins. The early ISC system in mitochondria – in addition to its role in mitochondrial Fe/S protein assembly – provides an unknown, sulfur- and possibly iron-containing substrate (X-S) to the cytosol via the ATP-dependent ABC transporter Atm1. The CIA system uses X-S for biogenesis of cytosolic and nuclear [4Fe-4S] proteins. How eukaryotes assemble cytosolic [2Fe-2S] proteins has not been clearly established and is in the focus of this work. In the bottom, known and potential cytosolic and nuclear [2Fe-2S] proteins are listed for *S. cerevisiae*, *H. sapiens*, and other eukaryotes (euk, also see Supplemental Table 2 in (49)). Cytosolic [2Fe-2S] proteins with a function in Fe/S protein assembly are given in bold.

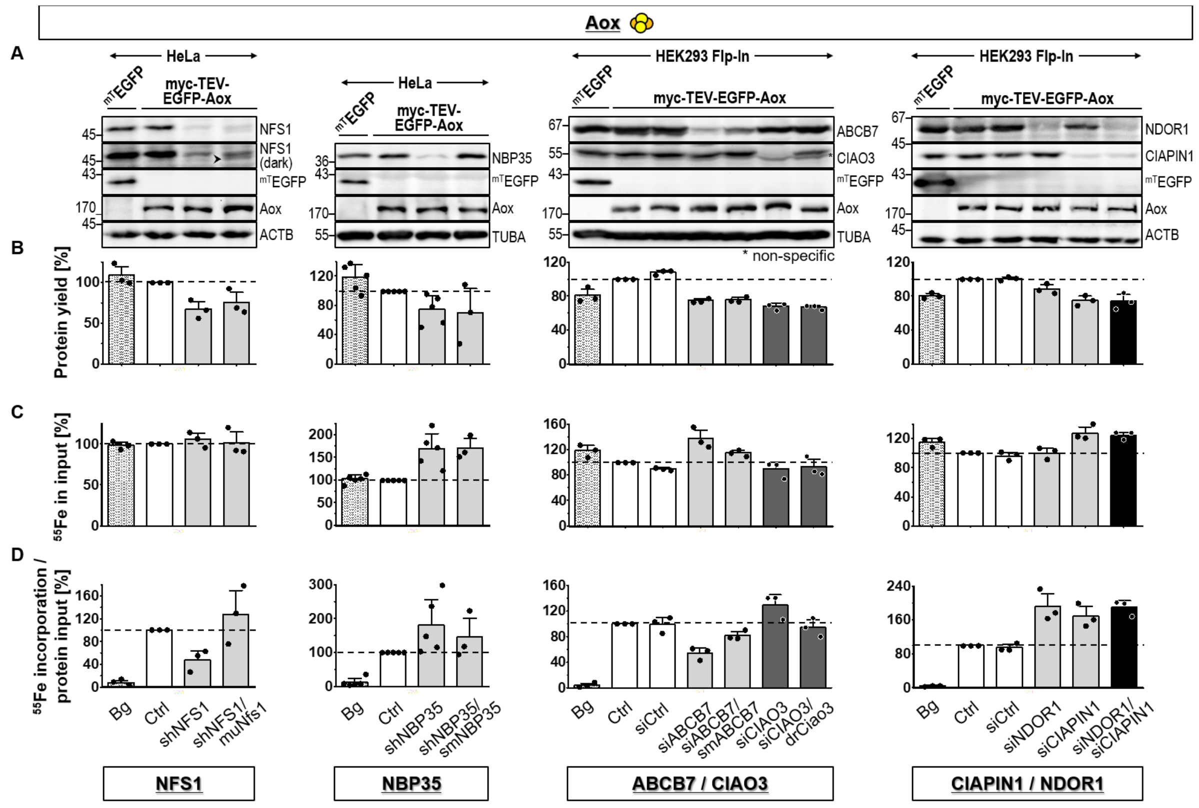

**Figure S2**: In relation to Figure 1A. [2Fe-2S] cluster assembly of aldehyde oxidase is dependent on ISC but not on CIA components. (**A**) HeLa or HEK293 Flp-In cell extracts from Fig. 1A or from cells transfected with control siRNA (siCtrl, sample 3 in panels 3 and 4) were subjected to immunoblotting against the indicated ISC and CIA factors, the reference myc-TEV-EGFP fusion protein, and the [2Fe-2S] reporter protein myc-TEV-EGFP-Aox (Aox). Beta-actin (ACTB) or alpha-tubulin (TUBA) served as loading controls. ABCB7 and CIAO3 depletion was performed in a conjoint experiment (third panel), and data is thus compared to a common set of control-transfected myc-TEV-EGFP (Bg) and myc-TEV-EGFP-Aox (Ctrl) samples (cf. Fig. 1A). The faint signal below the band for human NFS1 (first panel, arrowhead) represents expressed murine Nfs1 (cf. (35)). (**B**) Relative protein amount and (**C**) relative cellular $^{55}Fe$ content of input samples subjected to α-myc immunoprecipitation in Fig. 1A. (**D**) [2Fe-2S] incorporation into the myc-TEV-EGFP-Aox reporter protein was determined as the ratio of recovered $^{55}Fe$ radioactivity per input protein, and values (mean ±SD, $n \geq 3$) were normalized to those obtained for reporter protein-expressing control cells (Ctrl, set to 100%, white bars and dashed lines). Samples of cells transfected with control siRNAs (siCtrl) are also included into the data set. For abbreviations refer to Fig. 1.

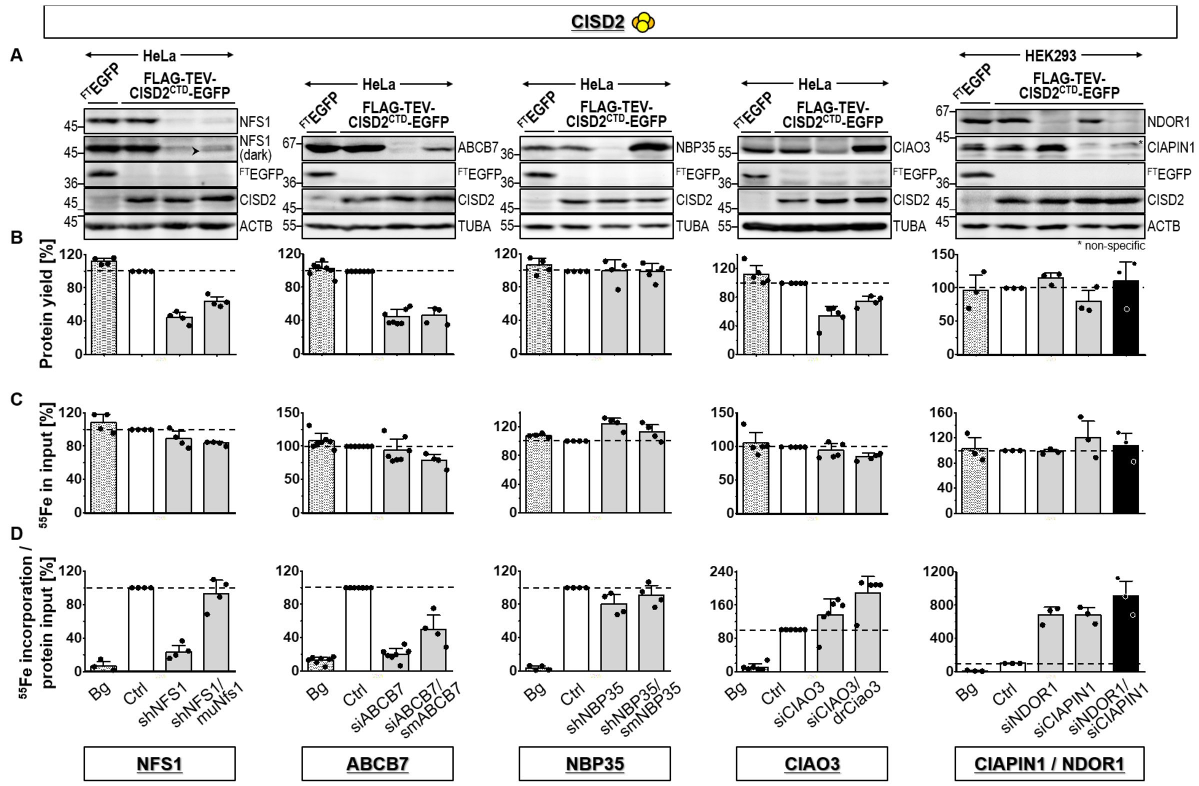

**Figure S3**: In relation to Figure 1B. [2Fe-2S] cluster assembly of CISD2 is dependent on ISC but not CIA components. (**A**) Aliquots of HeLa or HEK293 Flp-In cell extracts from Fig. 1B were subjected to immunoblotting against the indicated ISC and CIA factors, the reference FLAG-TEV-EGFP fusion protein, and the [2Fe-2S] reporter FLAG-TEV-$CISD2^{CTD}$-EGFP (CISD2). Beta-actin (ACTB) or alpha-tubulin (TUBA) served as loading controls. The faint signal below the band for human NFS1 (first panel, arrowhead) represents murine Nfs1 (cf. (35)). (**B**) Relative protein amount and (**C**) relative $^{55}Fe$ content of input samples subjected to α-FLAG immunoprecipitation in Fig. 1B. (**D**) [2Fe-2S] incorporation into the FLAG-TEV-$CISD2^{CTD}$-EGFP reporter protein was determined as the ratio of recovered $^{55}Fe$ radioactivity per input protein, and values (mean ± SD, n ≥ 3) were normalized to those obtained for reporter protein-expressing control cells (Ctrl, set to 100%, white bars and dashed lines). The FLAG-tagged EGFP fusion protein served as reference for non-specific $^{55}Fe$ background binding (Bg, first bars). For abbreviations refer to Fig. 1.

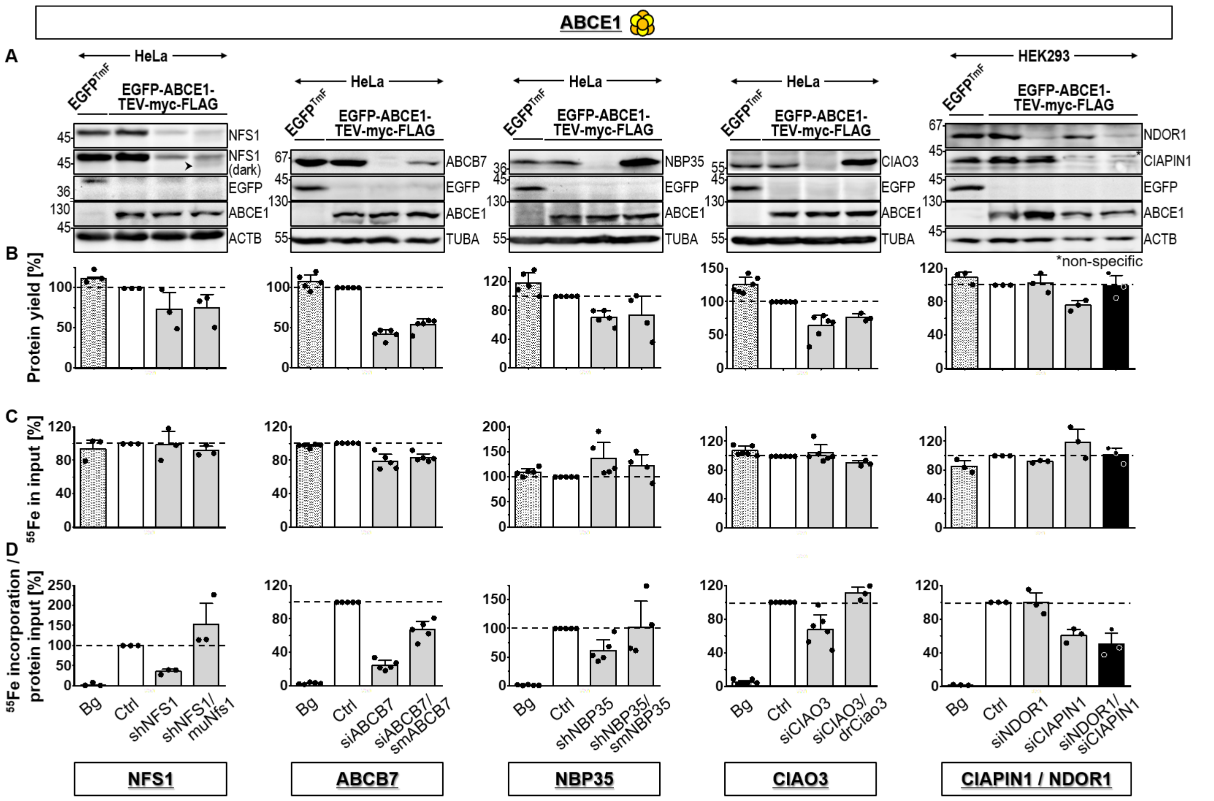

**Figure S4**: In relation to Figure 1C. [4Fe-4S] cluster assembly of ABCE1 is dependent on both ISC and CIA components. (**A**) Aliquots of HeLa or HEK293 Flp-In cell extracts from Fig. 1C were subjected to immunoblotting against the indicated ISC and CIA factors, the reference EGFP-TEV-myc-FLAG fusion protein (EGFP), and the [4Fe-4S] reporter EGFP-ABCE1-TEV-myc-FLAG (ABCE1). Beta-actin (ACTB) or alpha-tubulin (TUBA) served as loading controls. The faint signal below the band for human NFS1 (first panel, arrowhead) represents expressed murine Nfs1 (cf. (35)). (**B**) Relative protein amount and (**C**) relative $^{55}Fe$ content of input samples subjected to α-FLAG immunoprecipitation in Fig. 1C. (**D**) [4Fe-4S] incorporation into the EGFP-ABCE1-TEV-myc-FLAG reporter protein was determined as the ratio of recovered $^{55}Fe$ radioactivity per input protein, and values (mean ± SD, n ≥ 3) were normalized to those obtained for reporter protein-expressing control cells (Ctrl, set to 100%, white bars and dashed lines). The EGFP-TmF fusion protein served as reference for non-specific $^{55}Fe$ background binding (Bg). For abbreviations see Fig. 1.

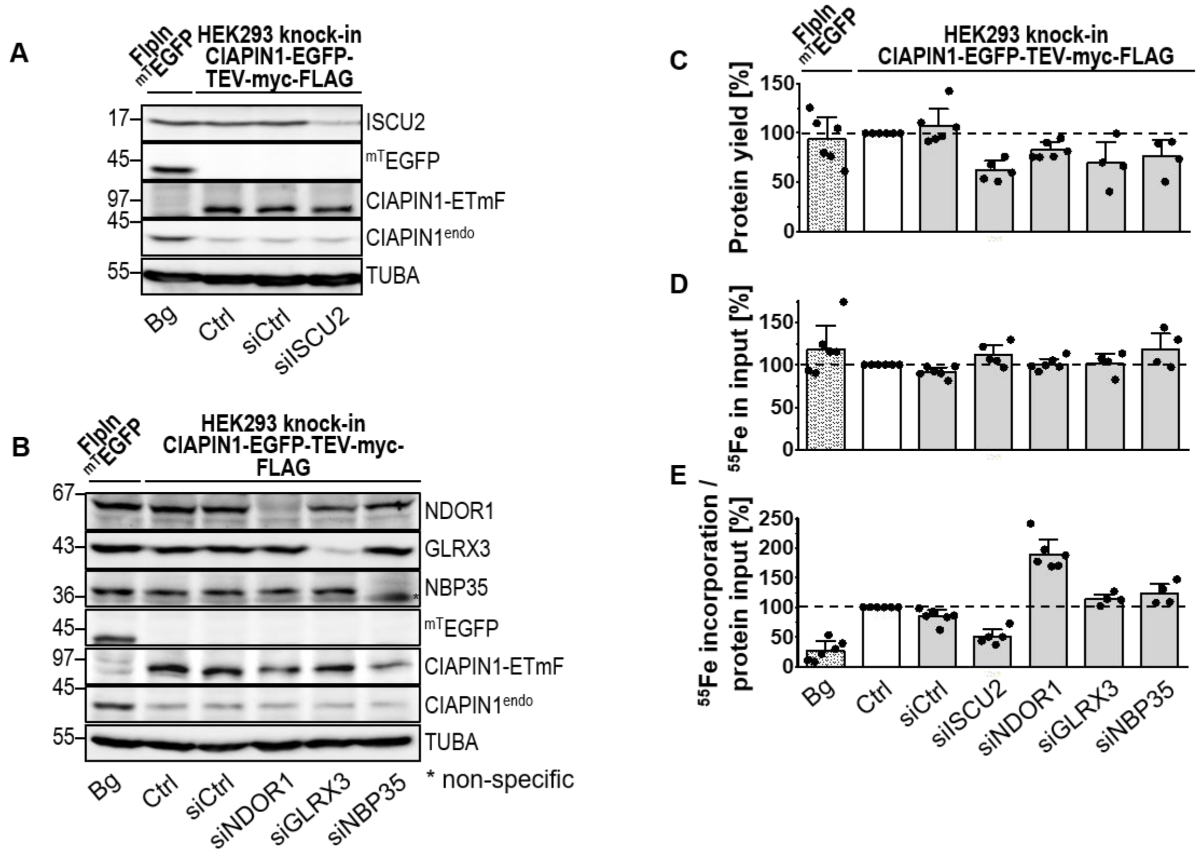

**Figure S5**: In relation to Figure 2A. Maturation of human CIAPIN1 is dependent on ISC but not CIA components. (**A,B**) Extracts from HEK293 CIAPIN1-EGFP-TEV-myc-FLAG knock-in cells or of myc-TEV-EGFP-expressing reference cells from Fig. 2A were subjected to immunoblotting against the indicated antigens. Alpha-tubulin (TUBA) served as loading control. (**C**) Relative protein amount and (**D**) relative $^{55}Fe$ amount of input samples used for α-myc immunoprecipitations. (**E**) Fe/S cluster incorporation into indicated reporter fusion proteins was determined as the ratio of recovered $^{55}Fe$ radioactivity per input protein, and is presented relative to the fusion reporter protein-expressing control cells (Ctrl, set to 100%, white bars and dashed lines). The myc-tagged EGFP fusion protein served as reference for non-specific $^{55}Fe$ background binding (Bg). Samples of cells transfected with control siRNAs (siCtrl) are also included into the data set. Values are given as mean ± SD, n ≥ 3. For abbreviations see Figs. 1 and 2.

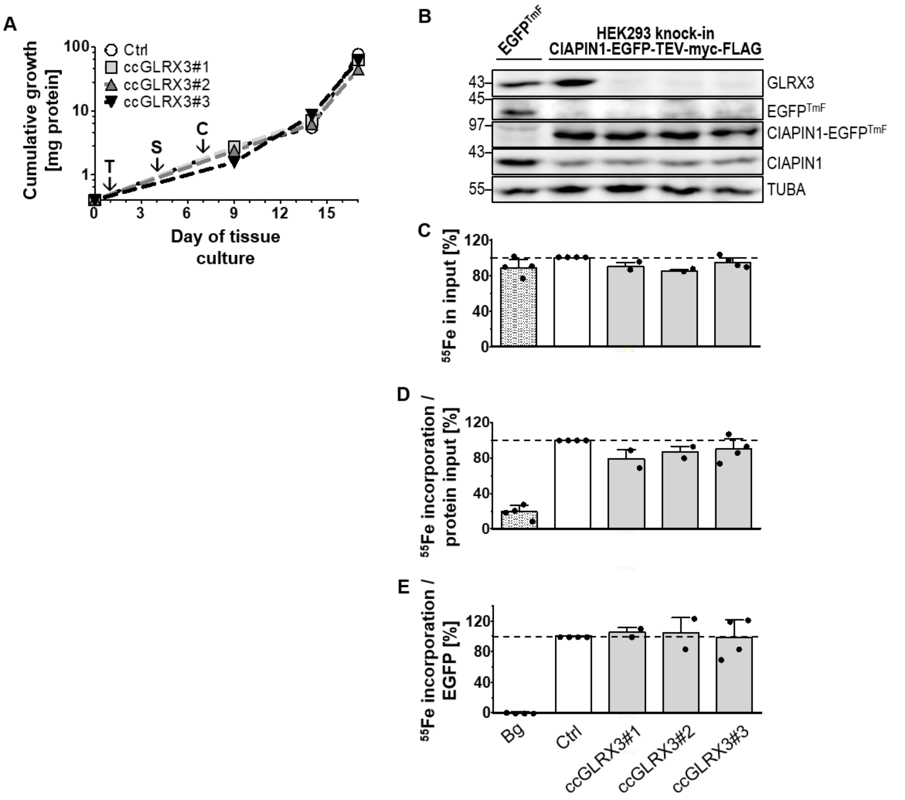

**Figure S6**: In relation to Figure 2B. CRISPR-Cas9-mediated *GLRX3* knock-out does not affect maturation of CIAPIN1. (**A**) HEK293 CIAPIN1-EGFP-TEV-myc-FLAG knock-in cells were seeded into culture vessels on day 0, transfected (T) on day 1 with one of three different *GLRX3*-directed guide RNAs (cc#1-3), and selected (S) by puromycin addition for 3 days. Cumulative growth of individual GLRX3 knock-out lines (cc#1, cc#2, cc#3) and of a control-transfected line (Ctrl) was estimated from the protein yield determined at various harvesting time points after puromycin removal (C, chase). (**B-E**) Cells from part A and EGFP-TEV-myc-FLAG-expressing cells (Bg; see below) were sub-cultured in presence of $^{55}$Fe-Tf for 3 to 4 days before $^{55}$Fe/S cluster maturation of CIAPIN1 was analyzed as in Fig. 2B (total GLRX3 depletion time 13 or 17 days). (**B**) Cell extracts were analyzed by immunostaining of the indicated proteins including the loading control alpha-tubulin (TUBA). (**C**) The relative $^{55}$Fe concentration in cell extracts used for $\alpha$-myc immunoprecipitation was determined by scintillation counting. (**D-E**) $^{55}$Fe/S cluster incorporation into CIAPIN1-EGFP was estimated as the ratio of co-precipitated $^{55}$Fe radioactivity either per recovered EGFP fluorescence (D) or per input protein (E). Transient expression of EGFP-TEV-myc-FLAG (EGFP$^{TmF}$) served as reference for non-specific background (Bg) co-precipitation of $^{55}$Fe. Values are presented relative to CIAPIN1-EGFP-TEV-myc-FLAG knock-in control cells (Ctrl, set to 100%, white bars and dashed lines) and are given as mean ± SD, n = 2 (ccGLRX3#1 and #2), n = 4 (ccGLRX3#3). For abbreviations see Figs. 1 and 2.

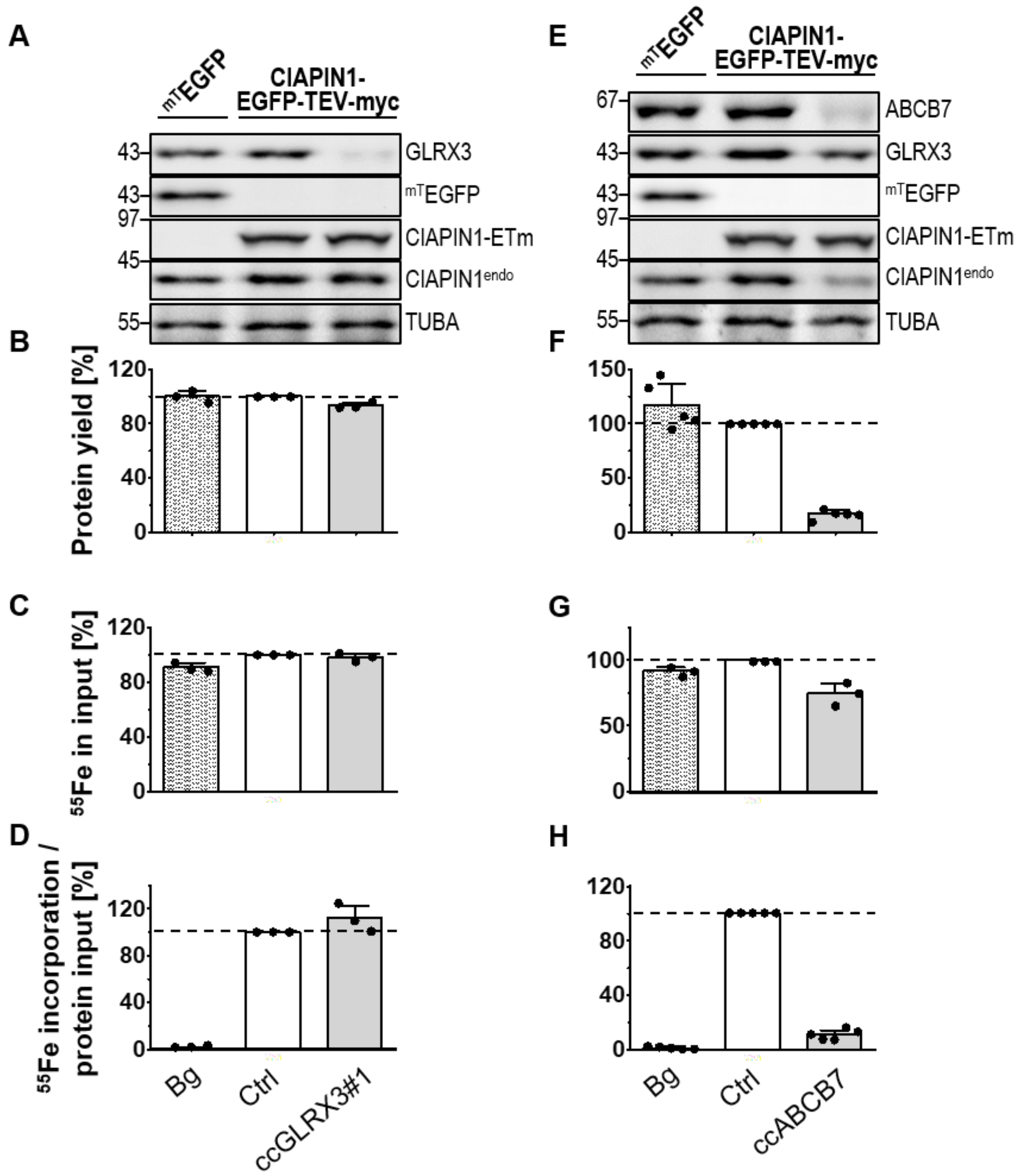

**Figure S7**: In relation to Figure 2C,D. Maturation of CIAPIN1 is dependent on ABCB7 but not on GLRX3. (**A,E**) Aliquots of HEK293 Flp-In cell extracts from Fig. 2C,D were subjected to immunoblotting against GLRX3, ABCB7, CIAPIN1, and the reference myc-TEV-EGFP fusion protein as indicated. Alpha-tubulin (TUBA) served as loading control. (**B,F**) Relative protein amount and (**C,G**) relative $^{55}$Fe content in cell extracts used for α-myc or α-FLAG immunoprecipitation. (**D,H**) $^{55}$Fe/S cluster incorporation into the indicated reporter proteins was determined as the ratio of immunoprecipitated $^{55}$Fe radioactivity per input protein, and is presented relative to the fusion reporter protein-expressing control cells (Ctrl, set to 100%, white bars and dashed lines). The myc-tagged EGFP fusion protein served as reference for non-specific $^{55}$Fe background binding (Bg). Values are given as mean ± SD, $n \geq 3$. For abbreviations see Figs. 1 and 2.

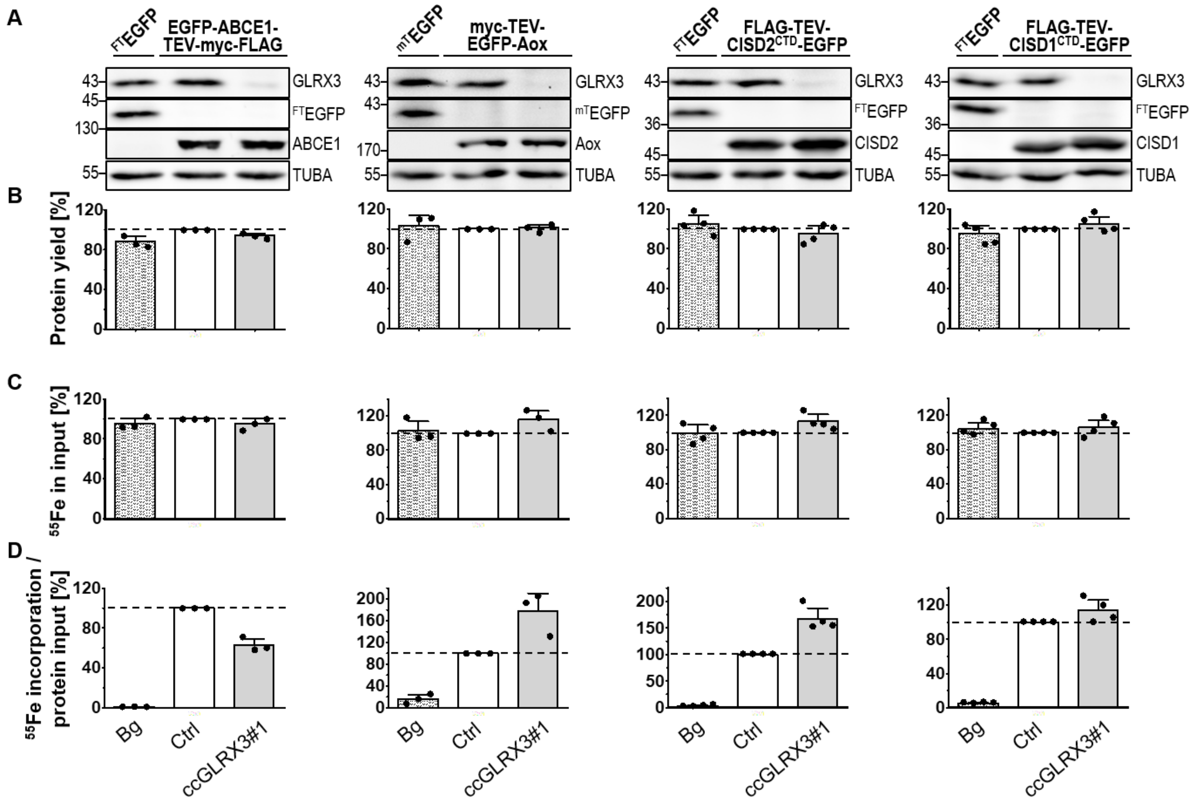

**Figure S8**: In relation to Figure 2E-H. Cytosolic [2Fe-2S] reporter proteins in human cells are matured independently of GLRX3. (**A**) Aliquots of HEK293 Flp-In cell extracts from Fig. 2E-H were subjected to immunoblotting against GLRX3 as well as Fe/S reporter or EGFP reference fusion proteins as indicated. Alpha-tubulin (TUBA) served as loading control. (**B**) Relative protein amount and (**C**) relative $^{55}$Fe content in cell extracts subjected to anti-myc or anti-FLAG immunoprecipitation. (**D**) $^{55}$Fe/S cluster incorporation into indicated reporter fusion proteins was determined as the ratio of immunoprecipitated $^{55}$Fe radioactivity per input protein, and is presented relative to the reporter protein-expressing control cells (Ctrl, set to 100%, white bars and dashed lines). FLAG- and myc-tagged EGFP fusion proteins served as reference for non-specific $^{55}$Fe background binding (Bg). Values are given as mean ± SD, $n \geq 3$. For abbreviations refer to Figs. 1 and 2.

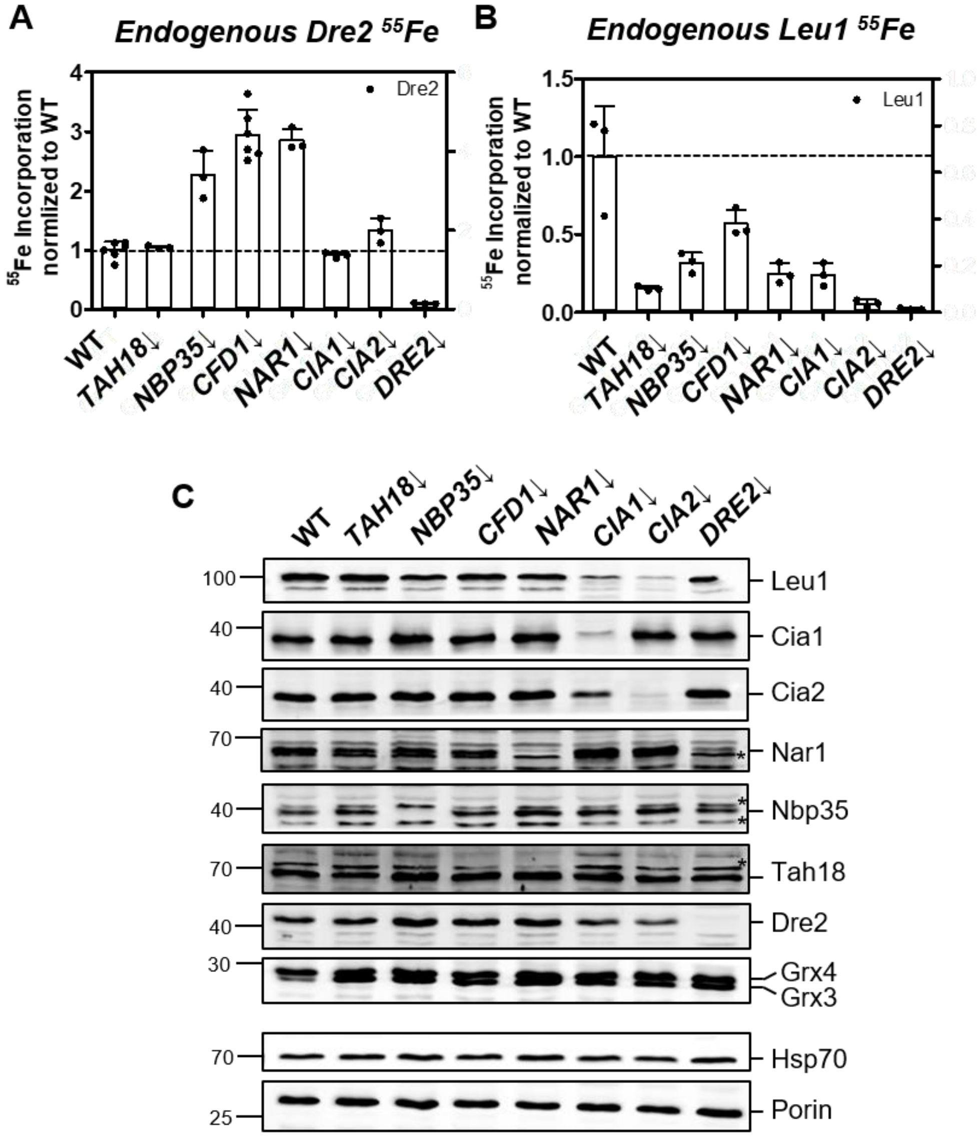

**Figure S9:** Endogenous yeast Dre2 does not require other CIA components for iron binding. (**A-B**) $^{55}$Fe radiolabeling experiments with endogenous Dre2 were performed due to the differences in protein levels observed between endogenous Dre2 (this study, Fig. 3D,F) and the previously employed Dre2-TAP construct in Gal-CIA strains (1), in which only in the former case lower protein levels were observed. WT and the indicated glucose-depleted (↓) *GAL* promoter-exchange CIA yeast strains were subjected to the $^{55}$Fe radiolabeling experiment as done in Fig. 3D-E. The CIA independence of Dre2 $^{55}$Fe binding is consistent with previous experiments utilizing the Dre2-TAP construct (31). Relative amounts of $^{55}$Fe bound to Dre2 and Leu1 were normalized to WT, where WT levels were $8.5 \pm 3.0 \times 10^3$ and $33.0 \pm 1.0 \times 10^3$ cpm/g cells, respectively. (**C**) Western blotting confirmed the depletion of CIA components in the indicated Gal-CIA strains. Values in bar charts represent the mean ± SD, n ≥ 3. * denotes non-specific bands observed in Western blotting.

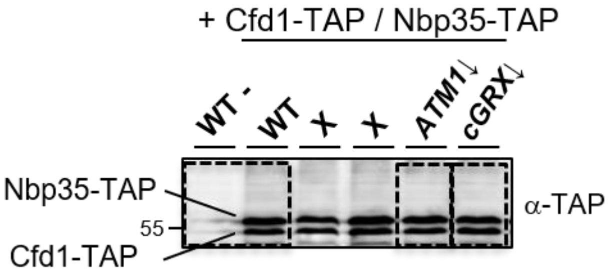

**Figure S10:** In relation to Figure 4B. Full Western blot image showing the anti-TAP staining for Cfd1-TAP and Nbp35-TAP in WT yeast cells and glucose-depleted Gal-*ATM1* and Gal-*GRX4grx3Δ* cells from Fig. 4B. Marked boxes indicate areas of the blots used. X denotes non-relevant lanes.

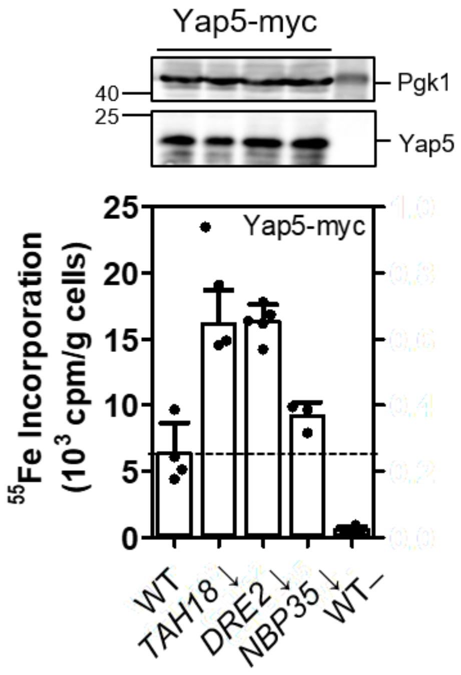

**Figure S11:** [2Fe-2S] cluster insertion into the reporter protein Yap5 is CIA-independent. WT and indicated (CIA protein-depleted) yeast strains were radiolabeled with $^{55}FeCl_3$ for 2 h followed by cell lysis (cf. Fig. 3C-E). Yap5-myc was immunoprecipitated using anti-myc beads, and $^{55}$Fe binding was quantified by scintillation counting. Protein expression was visualized by Western blotting (representative blots). Cytosolic Pgk1 served as loading control. WT cells transformed with empty plasmids are depicted as WT–. Values in bar charts represent the mean ± SD, $n \geq 3$.

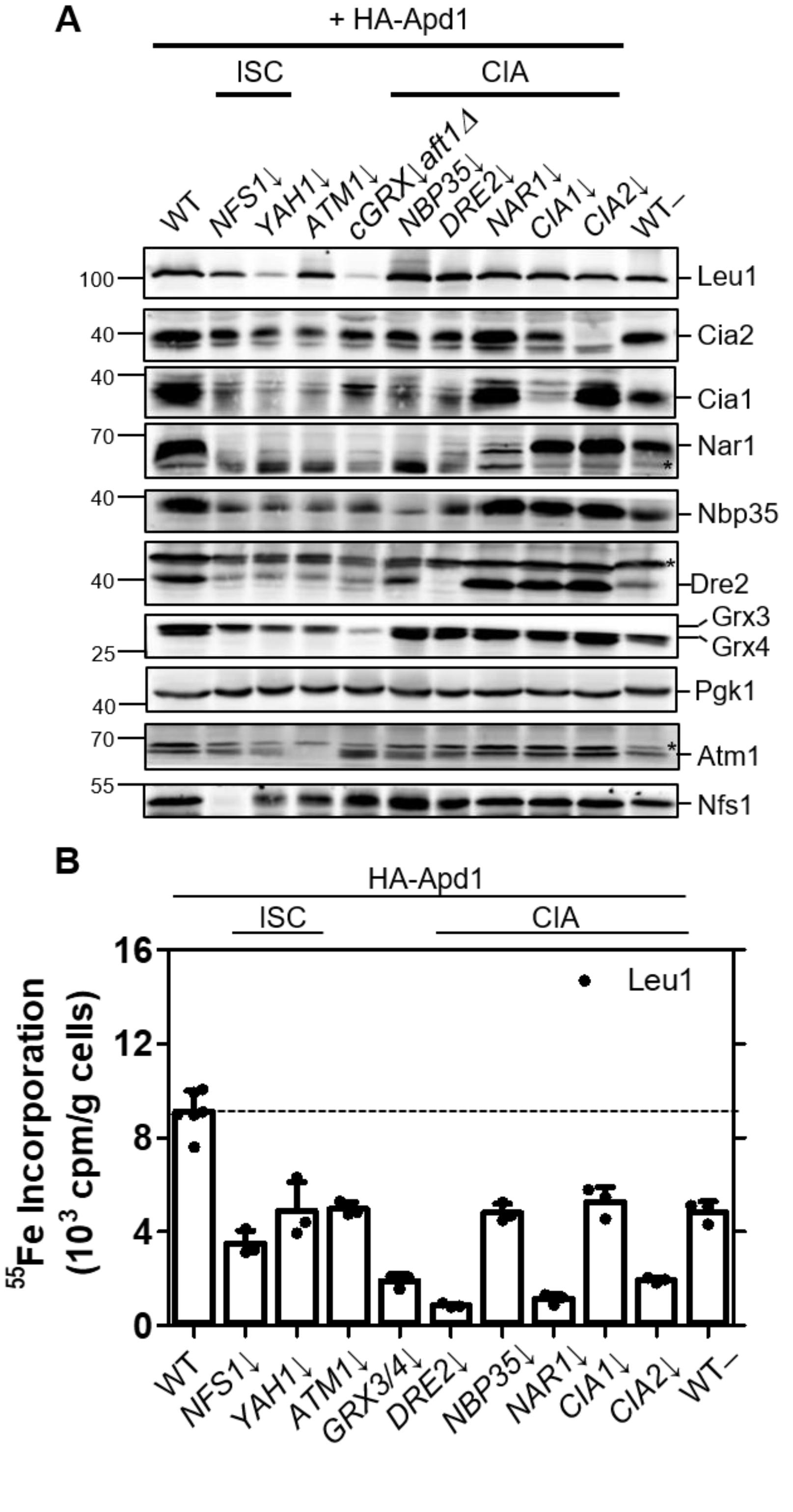

**Figure S12:** In relation to Figure 5. Like Apd1 (Fig. 5C), Leu1 depends on the ISC and CIA components for $^{55}$Fe/S cluster insertion. (**A**) Representative Western blots of extracts from wild-type (WT; W303) and the indicated ISC/CIA protein- or Atm1-depleted (↓) cells derived from $^{55}$Fe radiolabeling experiments of Fig. 5C. The Pgk1 loading control stain is the same as in Fig. 5C. The low-copy expression of HA-Apd1 caused a strong decrease in Yap5 staining in lanes 2-7 for unknown reasons. This behavior was not observed with other HA-Apd1 constructs

(Figs. S14C and S15A). Molecular mass markers are indicated in kDa. (**B**) After $^{55}Fe$ radiolabeling of the indicated cells, endogenous Leu1 was immunoprecipitated using specific antibodies. The horizontal dashed line indicates WT levels of $^{55}Fe$ binding. The control samples for WT cells transformed with an empty plasmid are depicted as WT−. Values represent the mean ± SD, n = 3. *, the asterisks denote non-specific bands observed in Western blotting in panels.

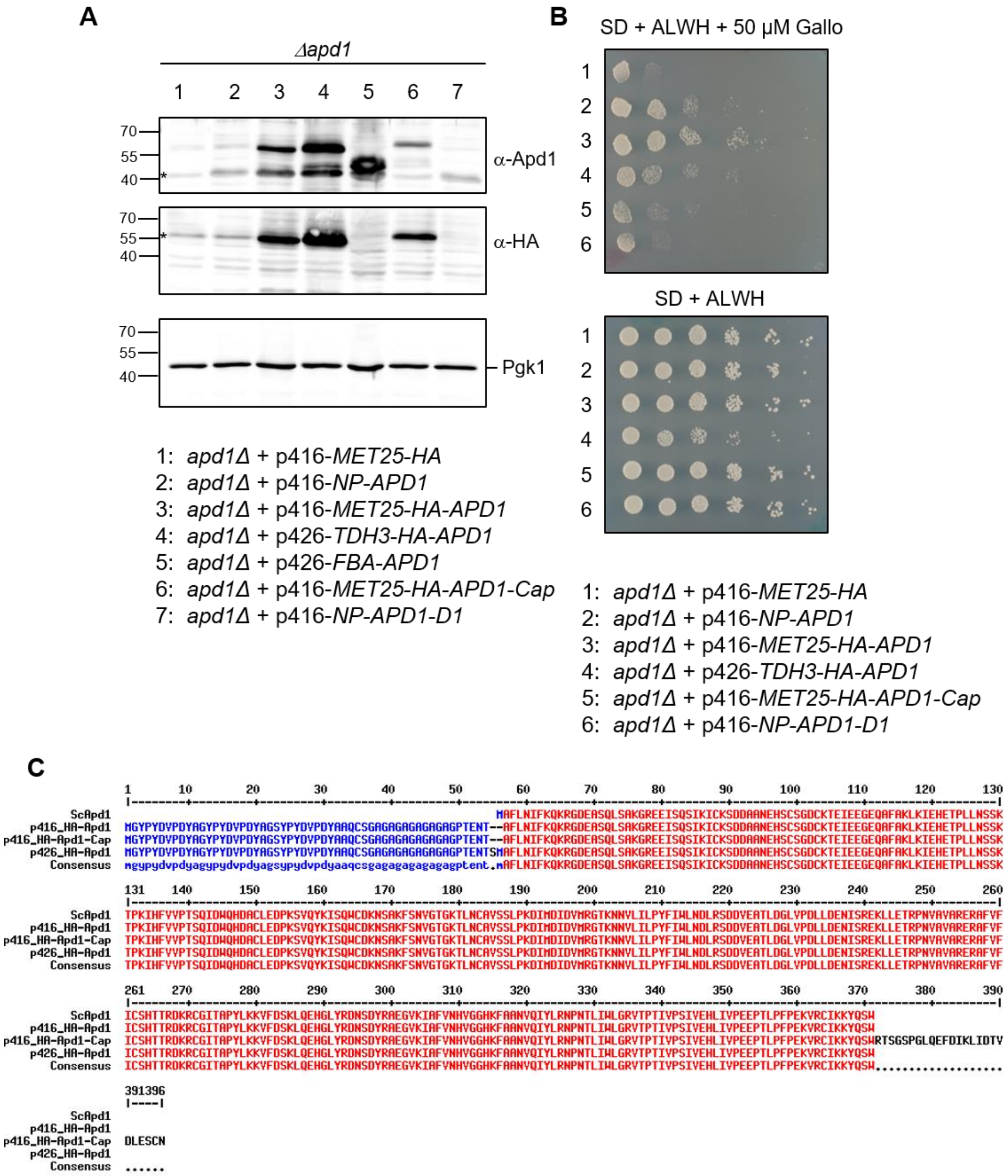

**Figure S13:** Expression level and the C-terminus of Apd1 influence its *in vivo* growth behavior. (**A-B**) The *apd1Δ* mutant strain was transformed with the indicated low- or high-copy plasmids coding for either wild-type or indicated mutant Apd1 proteins under the control of various promoters (MET25, low-copy; TDH3, high-copy; NP, natural promoter of *APD1*; or fructose 1,6-bisphostphate aldolase (FBA), high-copy -- refer to Table S4A) (19). Cells were grown in an overnight culture containing SC media, glucose (SD) and required supplements (A, adenine; L, leucine; W, tryptophan; H, histidine). On the following day, a fraction of the cells was used for a spot assay (B) or lysed for Western blot analysis (A). Protein levels of Apd1 under the control of its endogenous promotor (theoretical 35.8 kDa, lane 2) and overexpressed wild-type and mutant Apd1 proteins in cell lysates were assessed by Western blotting (A). Membranes were stained against endogenous Apd1 (top), the HA tag (middle), and Pgk1 as a loading control (bottom). *, the asterisks denote non-specific bands observed in Western blotting. (B) Serial dilutions (1:5) of the overnight cultures of *apd1Δ* containing the indicated plasmids were plated on SD agar plates containing required supplements and 50 μM gallobenzophenone (Gallo, top panel) followed by incubation for 3 days at 30°C. A control plate containing no Gallo served as a growth control (lower panel). (**C**) Multi-sequence alignment of the HA-Apd1 proteins used in this study compared to the native protein sequence of *S. cerevisiae* Apd1 (ScApd1). Refer to Supplementary Table S4A for construct information.

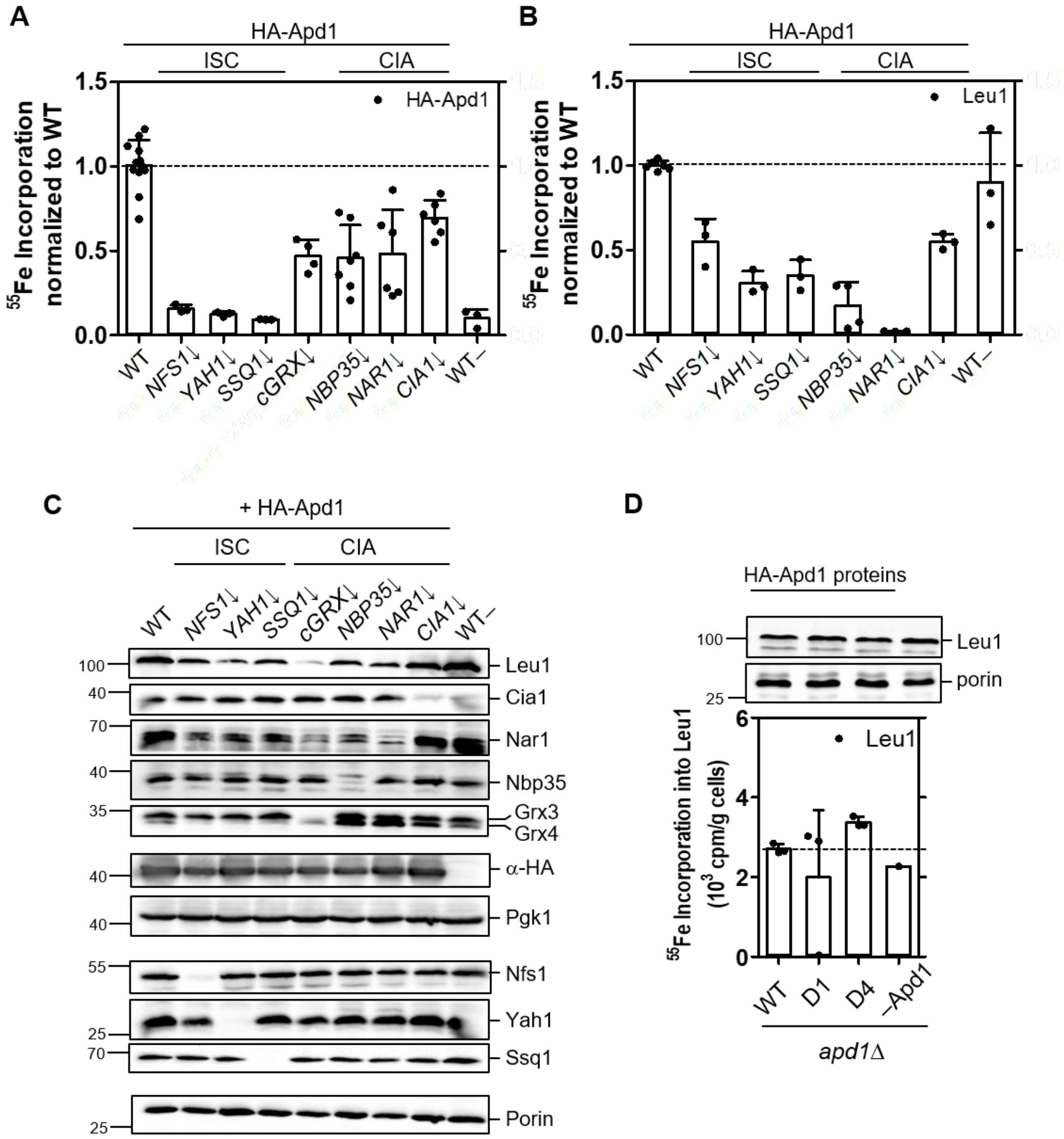

**Figure S14:** Overexpression of HA-Apd1 in wild-type (wild-type) and CIA protein-depletion strains from a yeast high-copy plasmid showed comparable results in $^{55}$Fe binding as low-copy expression. (**A-B**) $^{55}$Fe radiolabeling experiments utilizing the high-copy expression vector p426-*HA-APD1* in WT and ISC- or CIA-depleted (↓) yeast strains were performed as in Fig. 5C. $^{55}$Fe bound to immunoprecipitated HA-Apd1 (A) or endogenous Leu1 (B) was quantitated by scintillation counting. Bound $^{55}$Fe levels were normalized to the WT values ($14.5 \pm 6.6 \times 10^3$ and $3.8 \pm 1.1 \times 10^3$ cpm/g cells, respectively, depicted by dashed horizontal lines). The value for WT cells transformed with an empty plasmid as a control is depicted as WT–. (**C**) The expression of HA-Apd1 and the depletion efficiencies of ISC and CIA components were assessed by Western blotting. Porin and Pgk1 served as loading controls. Molecular mass markers are indicated in kDa. (**D**) As done in Fig. 5D, *apd1Δ* cells expressing WT or mutant

HA-tagged Apd1 proteins (D1, HA-Apd1-D1; D4, HA-Apd1-D4) from the high-copy vector were subjected to the $^{55}$Fe radiolabeling-immunoprecipitation assay and Leu1 was immunoprecipitated with homemade Leu1 antibodies. Values represent the mean ± SD, $n \geq 3$, (except for *apd1Δ* cells transformed with an empty plasmid (–Apd1) in D, where n=1).

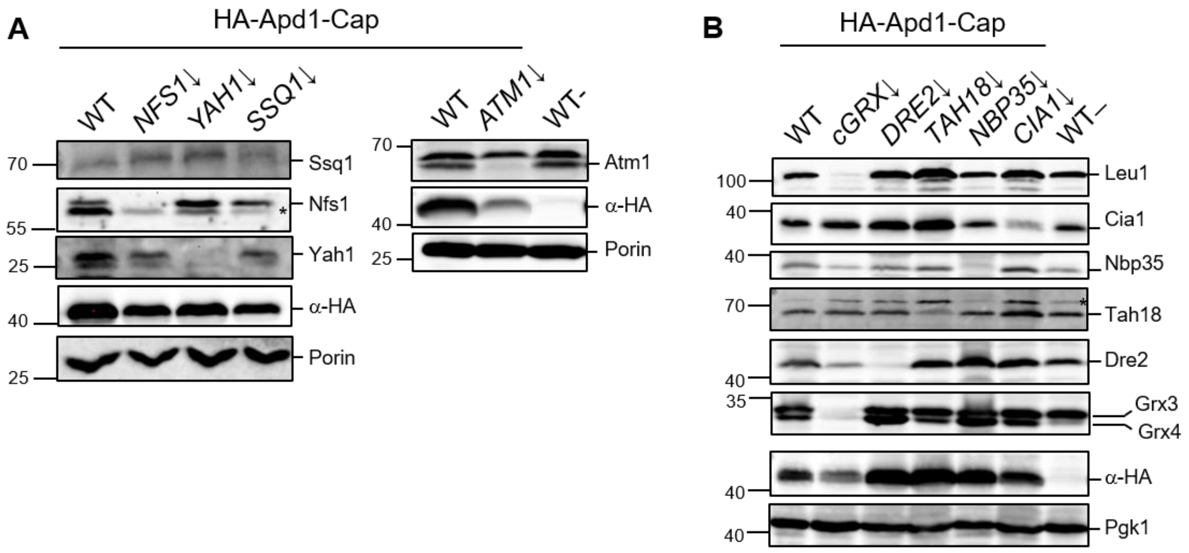

**Figure S15:** Western blot analysis for the $^{55}$Fe radiolabeling experiments shown in Figure 5E-F. (**A-B**) The overexpression of HA-Apd1-Cap and the depletion efficiencies of mitochondrial (C) and cytosolic Fe/S biogenesis components (D) were assessed by Western blotting. Molecular mass markers are indicated in kDa.

### Supplemental methods

#### *Cell culture and transfection*

HeLa and HEK293 cells were cultured in high-glucose DMEM (4.5 g/L) with standard supplementation of glutamine, penicillin-streptomycin, and fetal bovine serum (FBS). HEK293T FLP-IN T-REx cells (Invitrogen, (42)) were maintained in tetracycline-deprived FBS. pcDNA5/FRT/TO-derived plasmids (see below) were integrated into HEK293T FLP-IN T-REx cells according to the manufacturer's instructions, allowing for doxycycline-inducible (2 to 3 µg/mL) expression of EGFP-tagged Fe/S reporter constructs or of EGFP reference proteins. CIAPIN1 was genomically tagged with a EGFP-TEV-myc-FLAG fusion sequence via homology-directed repair (HDR, see below), achieved by co-transfection of a guide (g) RNA-encoding CRISPR plasmid directed against the 3' end of exon 8 (ccCIAPIN1, Table S4C), and the donor plasmid CIAPIN1(C)-EGFP-TEV-myc-FLAG-T2A-Kana#HDR (Table S4B). A stable cell line was established by G418 antibiotic selection.

Transfection of siRNAs and plasmids into HeLa cells by electroporation was carried out as described, using a Bio-Rad Gene Pulser Xcell system (40). HEK293 and HEK293T FLP-IN T-REx cells were transfected by a manually applied double pulse (225 V, 425 µF, exponential decay each) with a time interval of about 2 s. After transfection, HEK293 cells were seeded into collagen-coated flasks to improve cell adherence and recovery. Chemical transfection of CRISPR plasmids for gene knock-out (KO) in HEK293 and HEK293T FLP-IN T-REx cells was performed in collagen-coated culture vessels by the jetPRIME reagent (Polyplus, Illkirch, France) according to the manufacturer's instructions. Two to three days after transfection, KO cell lines were selected by puromycin addition (7.5 µg/mL) for 3-4 days, and analyzed within 3-4 weeks.

#### *RNAi, CRISPR, and plasmids*

*NFS1*- and *NBP35*-directed small hairpin (sh) RNA-encoding plasmids (Table S4D) as well as vectors for the transient expression of RNAi-resistant versions of the two proteins (shNFS1, shNBP35) were validated in previous studies (34, 35). *ABCB7*-directed and corresponding non-targeting (NoT) control (Ctrl) small interfering (si) RNAs were obtained from Horizon Discovery (Waterbeach, UK), whereas *CIAO3*, *NDOR1*, *CIAPIN1* genes (HGNC approved nomenclature, https://www.genenames.org/), and Silencer Select Negative Ctrl siRNA No.1 (Scr) were purchased from Ambion/ThermoFisher (Waltham, MA, USA). RNAi sequences are listed in Table S5A.

*ABCB7*-, *GLRX3*-, and *CIAPIN1*-directed gRNA sequences (Table S4C) for CRISPR-Cas9-mediated gene knock-out or knock-in in HEK293 cells were designed using sgRNA designer (Broad Institute (43)), E-CRISP (44), or the Guide Design Tool (http://crispr.mit.edu), respectively, and cloned into plasmid PX459 (Addgene, Watertown, MA, USA) according to reference (36).

Full-length human *ABCB7* (45) was cloned into the backbone of a pEGFP-series plasmid (Clontech), thereby replacing the EGFP sequence, and a subsequent PCR was performed according to Zheng et al. (46) to generate a silently mutated (sm), RNAi-resistant *ABCB7* version (mutagenesis primers are given in Table S5B). Zebrafish (*Danio rerio*, dr) *Ciao3* cDNA (clone IMAGp998J1016249Q) was obtained from Source Bioscience (Nottingham, UK), and the ORF was subcloned into the mammalian expression vector pExpress (47).

A murine aldehyde oxidase (Aox)-encoding plasmid was kindly provided by E. Garattini (Milan, Italy) and S. Leimkühler (Potsdam, Germany), and the Aox open reading frame (ORF) was subcloned into pEGFP-C1- (Clontech) and pcDNA5/FRT/TO- (Invitrogen) derived plasmids downstream of a myc-TEV-EGFP tag containing a tobacco edge virus (TEV) protease cleavage site. DNA sequences corresponding to the soluble C-terminal, [2Fe-2S] cluster-binding domains (CTD) of human CISD1 (aka mitoNEET, residues 33 to 108) or CISD2 (aka MINER1 or NAF-1, residues 56 to 135) with N-terminal FLAG-TEV-tags were cloned into plasmid pEGFP-N1, yielding FLAG-TEV-CISD2(CTD)-EGFP. The entire fusion sequences were subsequently subcloned into plasmid pcDNA5/FRT/TO. A reporter construct similar to EGFP-ABCE1-TEV-myc-FLAG has been described (40), and the fusion sequence was subcloned into plasmid pcDNA5/FRT/TO. The ORF of CIAPIN1 (31) was fused to a C-terminal EGFP-TEV-myc tag and cloned into pcDNA5/FRT/TO. pEGFP series or pcDNA5/FRT/TO backbones were used to clone reference plasmids by adding TEV-separated, N- or C-terminal myc- or FLAG-tag sequences to an EGFP ORF. Plasmids derived from the pcDNA5/FRT/TO backbone were subsequently integrated into HEK293T FLP-IN T-REx cells via pre-defined flippase recombinase (FLP) target (FRT) sites according to the FLP-IN T-Rex Core Kit guidelines (Invitrogen).

In order to genomically tag the 3' end of CIAPIN1 via homology-directed repair (HDR), a pEGFP-C1-based donor plasmid (CIAPIN1(C)-EGFP-TEV-myc-FLAG-T2A-Kana#HDR) was constructed including an EGFP-TEV-myc encoding sequence, a T2A peptide spacer, and a kanamycin resistance gene. The entire tripartite cassette was flanked by left and right homology arms derived from *CIAPIN1*, the length ranging from 900 to 1,000 bp, respectively

(Table S5C). Bacterial plasmid propagation and selection was achieved by exchanging the pUC origin of replication (ori) by a combined ColE1 ori-ampicillin resistance cassette obtained from the pMACS Kk.II plasmid via PciI and SbfI restriction sites. A summary of plasmids used in mammalian tissue culture is given in Tables S4B-D.

*$^{55}$Fe radiolabeling-immunoprecipitation assay*

Cells were cultured in the presence of $^{55}$Fe-loaded transferrin for up to three days, and reporter proteins were immunoprecipitated using anti-myc or anti-FLAG beads (48). After TEV protease-induced release of EGFP-tagged proteins from the affinity matrix, the eluate was analyzed for EGFP fluorescence as a measure for reporter protein recovery, and for $^{55}$Fe radioactivity as a measure for [2Fe-2S] and [4Fe-4S] cluster assembly. An EGFP-only reporter construct was included in order to monitor non-specific $^{55}$Fe background binding (Bg) which, however, was negligible.

**Tables**

**Table S1:** Nomenclature for yeast and human proteins involved in cellular Fe/S protein assembly

| **Mitochondrial ISC machinery** | | | |
|---|---|---|---|
| **Yeast** | **Human** | **Human (alternative)** | **Explanation** |
| Nfs1 | NFS1 | | NFS1 cysteine desulfurase; Nitrogen fixation 1-like protein |
| Isd11 | ISD11 | LYRM4 | LYR motif-containing 4 protein |
| Acp1 | ACP1 | NDUFAB1 | Acyl carrier protein |
| Isu1, Isu2 | ISCU2 | ISCU | Iron-sulfur cluster assembly enzyme |
| Yfh1 | FXN | Frataxin | Friedreich ataxia protein |
| Yah1 | FDX2 | FDX1L | Ferredoxin 2; Ferredoxin 1-like |
| Arh1 | FDXR | ADXR | Ferredoxin reductase; Adrenodoxin reductase |
| Ssq1 | HSPA9 | GRP75, PBP74, mortalin | Hsp70 chaperone |
| Jac1 | HSCB | HSC20, DNAJC20 | HscB iron-sulfur cluster co-chaperone homolog |
| Mge1 | GRPEL1,2 | hMGE | GrpE-like 1,2 |
| Grx5 | GLRX5 | GRX5 | Glutaredoxin 5 homologs |
| Isa1 | ISCA1 | ISA1, HBLD2 | HESB-like domain containing 2 |
| Isa2 | ISCA2 | ISA2, HBLD1 | HESB-like domain containing 1 |
| Iba57 | IBA57 | C1orf69 | Iron-sulfur cluster assembly factor of 57 kDa |
| Nfu1 | NFU1 | HIRIP5 | HIRA-interacting protein 5 |
| Bol1 | BOLA1 | - | BOLA family members 1 |
| Bol3 | BOLA3 | - | BOLA family members 3 |
| Ind1 * | NUBPL | IND1 | Nucleotide binding protein-like |
| *, not present in *S. cerevisiae* | | | |
| **Mitochondrial transporters** | | | |
| Atm1 | ABCB7 | ABC7 | ATP-binding cassette sub-family B member 7 |

| | | | |
|---|---|---|---|
| Mrs3, Mrs4 | MFRN1 | SLC25A37, mitoferrin 1 | Mitochondrial iron transporter 1 |
| Mrs3, Mrs4 | MFRN2 | SLC25A28, mitoferrin 2 | Mitochondrial iron transporter 2 |
| **CIA machinery** | | | |
| **Yeast** | **Human** | **Human (alternative)** | **Explanation** |
| Tah18 | NDOR1 | NR1; CIAE1 | NADPH dependent diflavin oxidoreductase 1 |
| Dre2 | CIAPIN1 | CIAE2; Anamorsin | Cytokine induced apoptosis inhibitor 1 |
| Nbp35 | NBP35 | NUBP1; CIAO5 | Nucleotide binding protein 1 |
| Cfd1 | CFD1 | NUBP2; CIAO6 | Nucleotide binding protein 2 |
| Nar1 | CIAO3 | NARFL; IOP1; HPRN | Nuclear prelamin A recognition factor-like;<br>Iron-only hydrogenase-like protein 1 |
| Cia1 | CIAO1 | CIAO1; CIA1; WDR39 | WD repeat domain 39 |
| Cia2 | CIAO2B | FAM96B; CIA2B; MIP18 | Family with sequence similarity 96 member B |
| -- | CIAO2A | FAM96A; CIA2A | Family with sequence similarity 96 member B |
| Mms19 (Met18) | MMS19 | MET18; CIAO4 | Methyl methanesulfonate sensitivity;<br>Methionine requiring |
| Lto1 | ORAOV1 | LTO1; TAOS1; CIAB1 | Oral cancer overexpressed protein 1A |
| Yae1 | YAE1 | YAE1D1; CIAB2 | Yae1 domain containing 1; "yet another essential gene" |
| **Proteins impacting CIA pathway (here demonstrated not to be essential)** | | | |
| Grx3-Grx4 | GLRX3 | PICOT | Glutaredoxin 3; Protein kinase C-interacting cousin of thioredoxin |
| Bol2[#] | BOLA2 | | BOLA family member 2 |
| [#], has no known impact on yeast CIA pathway | | | |

**Table S2:** Deletion phenotype of cytosolic monothiol glutaredoxins (cGrxs) studied to date.

| Organism | Protein | Strain/ Variant/ Cell type | Notes on depletion/deletion | Viable (Yes/No) | Reference |
|---|---|---|---|---|---|
| *Saccharomyces cerevisiae* | Grx3/4 | W303-1A | Constitutive iron-uptake response<br>Gal-*GRX4grx3Δaft1Δ* cells viable | N | (1); this study |
| *Saccharomyces cerevisiae* | Grx3/4 | BY 4742 | Hypoxia promotes growth, genetic suppressors identified | Y | (2, 3) |
| *Schizosaccharomyces pombe* | Grx4 | FY435 | Constitutive iron-uptake response | Y | (4) |
| *Aspergillus fumigatus* | GrxD | AfS77 | Grx domain essential<br>*ΔgrxD/ΔsreA* viable | N | (5) |
| *Cryptococcus neoformans* | Grx4 | H99 | Only thioredoxin domain essential | N | (6) |
| *Candida albicans* | Grx3 | HLY4492 | Constitutive iron-uptake response | Y | (7) |
| *Fusarium graminearum* | Grx4 | PH-1 | Decrease in iron-acquisition genes, increase in iron-consuming genes | Y | (8) |
| *Mus musculus* | GLRX3 | organismal | KO embryonically lethal | N | (9) |
| *Mus musculus* | GLRX3 | organismal | KO lethal at 12.5 days | N | (10) |
| *Mus musculus* | GLRX3 | Mammary epithelial cells<br>(cKO mice) | Conditional KO resulted in proliferation defects during alveolar development | Y | (11) |
| *Mus musculus* | GLRX3 | Cardio-myocytes<br>(cKO mice) | Conditional KO resulted in left ventricular hypertrophy at 12 months | Y | (12, 13) |
| *Arabidopsis thaliana* | GRXS17 | Col-0 | Minor effects on Fe/S proteins | Y | (14, 15) |
| *Homo sapiens* | GLRX3 | HEK293 | Late CIA defect, but not at CIAPIN1 | Y | This study |

**Table S3:** Yeast strains used in this study

| Strain | Genotype | Method of Generation | Depletion time | Source/ Reference |
|---|---|---|---|---|
| W303-1A | *MATα, ura3-1, ade2-1; trp1-1; his3-11,15; leu2-3,112* | - | - | (16) |
| YPH500 | *MAT*a, *ura3-52 lys2-801 ade2-101 trp1-63 his3-200 leu2-1* | - | - | (17) |
| *gsh1Δ* | YPH500; *gsh1::HIS3* | PCR fragment (pFA6a-*HIS3MX6*) | 48 h | (18) |
| *bol2Δ* | W3031-A, *bol2::NT2* | PCR fragment (pFA6a-*natNT2*) | - | This study |
| *aft1Δ* | W3031-A; *aft1::HIS3* | PCR fragment (pFA6a-*HIS3MX6*) | - | This study |
| *apd1Δ* | W3031-A, *apd1::NT2* | PCR fragment (pFA6a-*natNT2*) | - | (19) |
| Gal-*NFS1* | W303-1A; *pNFS1::GAL1-10-HIS3* | PCR fragment ( ) | 40 h | (20) |
| Gal-*YAH1* | W303-1A; *pYAH1::GALL-natNT2* | PCR fragment (pYM-N24) | 40 h | (21) |
| Gal-*SSQ1* | W303-1A; *pSSQ1::GAL1-10-HIS3* | PCR fragment (pFA6a-*HIS3-Gal*) | 64 h | (20) |
| Gal-*ATM1* | W303-1A; *pATM1::GALL-natNT2* | PCR fragment (pYM-N24) | 64 h | (22) |
| Gal-*GRX4 grx3Δ* | W3031-1A; *pGRX4::GALL-natNT2; grx3::LEU2* | W303-1A *grx3Δ*; PCR fragments (pYM-N27) (23) | 48 h | (1) |
| Gal-*GRX4 grx3Δaft1Δ* | W3031-1A; *pGRX4::GALL-natNT2; grx3::LEU2, aft1::HIS3* | W303-1A Gal-*GRX4 grx3Δ*,PCR fragment (pFA6a-*HIS3MX6*) | 48 h | This study |
| Gal-*DRE2* | W303-1A; *pDRE2::GAL1-10-natNT2* | PCR fragment (pYM-N23) | 16 h | (1) |
| Gal-*TAH18* | W303-1A; *pTAH18::GAL1-10-natNT2* | PCR fragment (pYM-N23) | 72 h | This study |
| Gal-*NBP35* | W303-1A; *pNBP35::GAL1-10-HIS3* | PCR fragment (pFA6a-*HIS3-Gal*) | 40 h | (24) |
| Gal-*CFD1* | W303-1A; *pCFD1::GAL1-10-HIS3* | PCR fragment (pFA6a-*HIS3-Gal*) | 64 h | (25) |
| Gal-*NAR1* | W303-1A; *pNAR1::GAL1-10-HIS3* | PCR fragment (pFA6a-*HIS3-Gal*) | 40 h | (26) |
| Gal-*CIA2* | W303-1A; *pCIA2::GAL1-10-natNT2* | PCR fragment (pFA6a-*natNT2-Gal*) | 40 h | (27) |
| Gal-*CIA1* | W303-1A; *pCIA1::GAL1-10-HIS3* | PCR fragment (pFA6a-*HIS3-Gal*) | 40 h | (25) |

**Table S4A:** Plasmid constructs used for analyses in yeast

| Plasmid | Figure | ORF | Backbone | Source/Reference |
|---|---|---|---|---|
| p426-*YAP5-AD-myc* | 6D, 7B, S11 | *YAP5* actuator domain Δ1-E115, C-terminal myc | p426-*TDH3* | (28) |
| p416-*NP-APD1* | S13A,B | *APD1* | p416 | (19) |
| p416-*NP-APD1-D1* | S13A,B | *APD1*, C-terminal W removed | p416 | (19) |
| p416-*3HA-APD1* | 5C, 6C, S12, S13 | *APD1*, N-terminal 3HA | p416-*MET25* | This study |
| p416-*3HA-APD1-Cap* | 5E-F, S13, S15 | *APD1*, N-terminal 3HA + 25 residue C-terminal extension | p416-*MET25* | This study |
| p426-*3HA-APD1* | 5D, 7A, S13, S14 | *APD1*, N-terminal 3HA | p426-*TDH3* | This study |
| p426-*3HA-APD1-D1* | 5D | *APD1* | p416 | This study |
| p426-*3HA-APD1-D4* | 5D | *APD1* | p416 | This study |
| p426-*FBA-APD1* | S13A | *APD1* | p426 | (19) |
| p424-*BOL2-HA* | 6A-B | *BOL2*, C-terminal 3HA | p426-*TDH3* | This study |
| p424-*GRX4-myc* | 6E | *GRX4*, C-terminal myc | p424-*TDH3* | This study |
| p426-*NAR1-3HA* | 6H | *NAR1*, C-terminal 3HA | p426-*TDH3* | (26) |
| p416-*NAR1-3HA-* | 7D | *NAR1*, N-terminal 3HA | p416-*MET25* | (26) |
| p426-*NBP35-TAP* | 4A-D, 6G, S10 | *NBP35*, C-terminal TAP | p426-*TDH3* | (24) |
| p414-*NBP35-TAP* | 4E, 7E | *NBP35*, C-terminal TAP | p416-*MET25* | This study |
| p424-*CFD1-TAP* | 4A-D, S10 | *CFD1*, C-terminal TAP | p424-*TDH3* | (29) |
| p416-*3HA-CFD1* | 4E, 7E | *CFD1*, N-terminal 3HA | p416-*MET25* | (30) |
| p424-*DRE2-TAP* | 6F | *DRE2*, C-terminal TAP | p424-*TDH3* | (31) |
| p424-*RAD3-TAP* | 4A-B | *RAD3*, C-terminal TAP | p424-*TDH3* | (32) |
| p426-*ILV3-3HA* | 7C | *ILV3*, C-terminal 3HA | p426-*TDH3* | (1) |
| pFET3-*GFP* | 3B | *GFP; FET3* promoter replacing *MET25* promoter | p416-*MET25* | (33) |

**Table S4B:** Plasmid constructs used for protein overproduction in mammalian cells

| Plasmid | Figure | ORF | Backbone | Source/Reference |
|---|---|---|---|---|
| pMCS_smABCB7 | 1, S2-4 | *human ABCB7*, silently mutated | pEGFP series (Clontech) | This study |
| pMCS_EGFP-ABCE1-TEV-myc-FLAG | 1C, S4 | *human ABCE1*, N-terminal EGFP, C-terminal TEV-myc-FLAG | pEGFP series (Clontech) | This study |
| pcDNA5_EGFP-ABCE1-TEV-myc-FLAG | 2E, S8 | *human ABCE1*, N-terminal EGFP, C-terminal TEV-myc-FLAG | pcDNA5/FRT/TO (Invitrogen) | This study |
| pMCS_myc-TEV-EGFP-Aox | 1A, S2 | *murine Aox*, N-terminal myc-TEV-EGFP | pEGFP series (Clontech) | This study |
| pcDNA5_myc-TEV-EGFP-Aox | 1A, 2F, S2, S8 | *murine Aox*, N-terminal myc-TEV-EGFP | pcDNA5/FRT/TO (Invitrogen) | This study |
| pEXPR_drCiao3 | 1, S2-4 | *Danio rerio* (dr) *Ciao3* | pExpress | This study |
| pcDNA5_CIAPIN1-EGFP-TEV-myc | 2C-D, S7 | *human CIAPIN1*, C-terminal EGFP-TEV-myc | pcDNA5/FRT/TO (Invitrogen) | This study |
| CIAPIN1(C)-EGFP-TEV-myc-FLAG-T2A-Kana#HDR | 2A-B, S5-6 | *human CIAPIN1 (aa277-312)*, C-terminal EGFP-TEV-myc-FLAG | pEGFP series (Clontech) | This study |
| pcDNA5_FLAG-TEV-CISD1(CTD)-EGFP | 2H, S8 | *human CISD1* (Δ 1-32), N-terminal FLAG-TEV, C-terminal EGFP | pcDNA5/FRT/TO (Invitrogen) | This study |
| pMCS_FLAG-TEV-CISD2(CTD)-EGFP | 1B, S3 | *human CISD2* (Δ 1-55), N-terminal FLAG-TEV, C-terminal EGFP | pEGFP series (Clontech) | This study |
| pcDNA5_FLAG-TEV-CISD2(CTD)-EGFP | 2G, S8 | *human CISD2* (Δ 1-55), N-terminal FLAG-TEV, C-terminal EGFP | pcDNA5/FRT/TO (Invitrogen) | This study |
| pMCS_EGFP-TEV-myc-FLAG | 1C, 2B, S4, S6B-E | *EGFP*, C-terminal TEV-myc-FLAG | pEGFP series (Clontech) | This study |
| pMCS_myc-TEV-EGFP | 1A, S2 | *EGFP*, N-terminal myc-TEV | pEGFP series (Clontech) | This study |
| pcDNA5_myc-TEV-EGFP | 1A, 2A,C-D,F, S2, S5, S7-8 | *EGFP*, N-terminal myc-TEV | pcDNA5/FRT/TO (Invitrogen) | This study |
| pMCS_FLAG-TEV-EGFP | 1B, S3 | *EGFP*, N-terminal FLAG-TEV | pEGFP series (Clontech) | This study |
| pcDNA5_FLAG-TEV-EGFP | 2E,G-H, S8 | *EGFP*, N-terminal FLAG-TEV | pcDNA5/FRT/TO (Invitrogen) | This study |
| pMCS_smNBP35 | 1, S2-4 | human *NBP35*, silently mutated | pEGFP series (Clontech) | (34) |
| pMCS_muNfs1 | 1, S2-4 | murine (mu) *Nfs1* | pEGFP series (Clontech) | (35) |

**Table S4C:** CRISPR-related plasmid constructs used for analyses in mammalian cells

| Plasmid | Figure | gRNA target sequence | Backbone | Source/Reference |
|---|---|---|---|---|
| ccABCB7 | 2D, S7E-H | TCACAGTTGCAGTCACACGG | PX459 (36) | This study |
| ccCIAPIN1 | 2A-B, S5-6 | CAATCTTCATGATGCCTAGG | PX459 (36) | This study |
| ccGLRX3#1 | 2C,E-H, S6, S7A-D,S8 | GGCTGAGCCGACCTCCTCCA | PX459 (36) | This study |
| ccGLRX3#2 | S6 | GAGTCAATTTCTTCAAGCGA | PX459 (36) | This study |
| ccGLRX3#3 | 2B, S6 | GGACTCAAAGCCTATTCCAGT | PX459 (36) | This study |

**Table S4D:** shRNA plasmid constructs used for gene silencing in mammalian cells

| Plasmid | Figure | shRNA target sequence | Backbone | Source/Reference |
|---|---|---|---|---|
| shNFS1 | 1, S2-4 | GCUGAGGGCUUUCAGGUCAU | pSUPER (37) | huNFS1-R3 (35) |
| shNBP35 | 1, S2-4 | CGUUAGCCUACAGAAGUAU | pSuperior (OligoEngine) | (34) |

**Table S5A:** RNAi sequences

| Target | Figure | Source | siRNA ID | Target sequence |
|---|---|---|---|---|
| ABCB7 | 1, S2-4 | Horizon Discovery | J-007305-05 | UAGACUCACUGCUGAAUUA |
| | | Horizon Discovery | J-007305-08 | CAACAGCAGUUCUGAUUGG |
| CIAO3 | 1, S2-4 | Ambion/ThermoFisher | s34746 | AGAUUCGCAUUGAAGAUGA |
| | | Ambion/ThermoFisher | s34747 | AGACCGUGCUUAUCACCCA |
| | | Ambion/ThermoFisher | s34748 | AGAAGGUUCUAGAUGCUAA |
| CIAPIN1 | 1, S2-4 | Ambion/ThermoFisher | s32591 | GGUUCUUCUAGGCAGCUUA |
| | | Ambion/ThermoFisher | s32592 | GCAAUCUGCCCACAAAGAA |
| | | Ambion/ThermoFisher | s32593 | AGACAGCUGUAGAUAACAA |
| | | Ambion/ThermoFisher | s238467 | ACAGCUGUAGAUAACAAUA |
| | | Ambion/ThermoFisher | s238466 | AGUACAGUCUGUUCGAGAA |
| GLRX3 | 2A, S5B-E | Horizon Discovery | Custom (38) | GCCUAUUCCAGUUGGCCUA |
| ISCU2 | 2A, S5A,C-E | Horizon Discovery | J-012837-10 | GAGCUAUGAGAUACGCACA |
| | | Horizon Discovery | J-012837-11 | CAGCAUGUGGUGACGUAAU |
| NBP35 | 2A, S5B-E | Ambion/ThermoFisher | Custom (34) | CGUUAGCCUACAGAAGUAU |
| NDOR1 | 1, 2A, S2-4, S5B-E | Ambion/ThermoFisher | S25924 | GGAGUUCAGUUCUGCCCAA |
| | | Ambion/ThermoFisher | S25926 | GCUGUAGUGCAGUUCCAGA |
| NoT* | S2, S5 | Horizon Discovery | D-001810-01 | UGGUUUACAUGUCGACUAA |
| Scr* | S2, S5 | Ambion/ThermoFisher | 4390843 | not available |

*Used as a mixture for control siRNA (siCtrl) application.

**Table S5B:** PCR primers used for site-directed mutagenesis of human ABCB7

| siRNA | Forward primer (5' => 3') | Reverse primer (5' => 3') |
|---|---|---|
| J-007305-05 | ctactgctgtactcatcggctatggtgta tcaagagctggag | agccgatgagtacagcagtagccatggtt gcaactgtatttggt |
| J-007305-08 | tattgatagtctcttaaactatgaaactg tgaagtattttaataatgaaagatatgaa | catagtttaagagactatcaatagcagca ttacctgcatcattatc |

Silent mutations are indicated in red

**Table S5C:** PCR primers for amplification of restriction site-containing left and right homology arms to add a C terminal genomic EGFP-TEV-myc-FLAG-tag to CIAPIN1

| Homology arm (HA) | Forward primer (5' => 3') | Reverse primer (5' => 3') |
|---|---|---|
| Left (LHA) | CP1_LHA_**PciI** ta**ACATGT**GGGGATTAGACCTTCTTC TAAACTA | CP1_LHA_**SalI** tat**GTCGAC**GGCATCATGAAGATTGCTATCAC |
| Right (RHA) | CP1_RHA_**PacI** ac**TTAATTAA**TCTGCTCCTCCAGCCA | CP1_RHA_**SbfI** at**CCTGCAGG**TCTCTTAAATCCACACAATC |

Restriction sites are indicated in bold

**Table S6:** Antibodies used in this study

| Yeast | | | |
|---|---|---|---|
| **Antibody** | **Validation** | **Dilution used** | **Source (product #, RRID)** |
| rabbit anti-Cia1 | Lill laboratory, validated in house by regulatable expression of *CIA1* via a *Gal1-10* promoter | 1:1000 | (25) |
| rabbit anti-Cia2 | Lill laboratory, validated in house by regulatable expression of *CIA2* via a *Gal1-10* promoter | 1:1000 | (27) & this study |
| rabbit anti-Dre2 | Lill laboratory, validated in house by regulatable expression of *DRE2* via a *Gal1-10* promoter | 1:1000 | This study |
| rabbit anti-Grx3/4 | Lill laboratory, validated in house by regulatable expression of *GRX4* via a *GalL* promoter | 1:1000 | (1) |
| rabbit anti-Leu1 | Lill laboratory, validated in house by *LEU1* deletion strain (*leu1Δ*) | 1:1000 | (27) & this study |
| rabbit anti-Nar1 | Lill laboratory, validated in house by regulatable expression of *NAR1* via a *Gal1-10* promoter | 1:1000 | (26) |
| rabbit anti-Nbp35 | Lill laboratory, validated in house by regulatable expression of *NBP35* via a *Gal1-10* promoter | 1:1000 | (24) |
| rabbit anti-Nfs1 | Lill laboratory, validated in house by regulatable expression of *NFS1* via a *Gal1-10* promoter | 1: 1000 | (20) |
| mouse anti-Pgk1 | Validated by company | 1:1000 | Abcam (ab113687, RRID:AB_2161220) |
| rabbit anti-Por1 | Lill laboratory, R.L. | 1:1000 | - |
| rabbit anti-Ssq1 | Lill laboratory, validated in house by regulatable expression of *SSQ1* via a *Gal1-10* promoter | 1:1000 | (20) |
| rabbit anti-Tah18 | Lill laboratory, validated in house by regulatable expression of *TAH18* via a *Gal1-10* promoter | 1:1000 | This study |
| rabbit anti-Yah1 | Lill laboratory, validated in house by regulatable expression of *YAH1* via a *GalL* promoter | 1:1000 | (20) & this study |
| rabbit anti-Yap5 | Lill laboratory, validated in house by immunoprecipitation using yeast cells transformed with plasmids lacking Yap5 antigen | 1:1000 | (28) |
| rabbit anti-Apd1 | Pierik laboratory | 1: 1000 | (19) |

| Human | | | |
|---|---|---|---|
| **Antibody** | **Validation** | **Dilution used** | **Source (product #)** |
| rabbit anti-ABCB7 | Validated in house by RNAi (this study) | 1:1000 | St. John's Laboratory (STJ116899) |

| | | | |
|---|---|---|---|
| rabbit anti-ABCE1 | Validated by RNAi and immunoprecipitation (personal observation and this study) | 1:1000 (affinity purified) | K. Sipos (Pecs, Hungary) |
| mouse anti-beta actin (clone C4) | validated by the manufacturer and by Bräutigam et al. (39) | 1:1000 | Santa Cruz Technology (sc-47778, RRID:AB_2714189) |
| rabbit anti-aldehyde oxidase | Validated by immunoprecipitation (this study) | 1:666 (plus ABC) | Abcam (ab197828, RRID:AB_2920908) |
| rabbit anti-CIAO3 | Lill laboratory, validated in house by immunoprecipitation and RNAi (e. g. this study) | 1:25 (affinity purified) | (40) |
| rabbit anti-CIAPIN1 | Lill laboratory, validated in house by RNAi (this study) | 1:5000 | This study |
| rabbit anti-GFP | Lill laboratory, validated in house by overexpression | 1:300 (purified) | This study |
| rabbit anti-GLRX3 | validated in house by RNAi and CRISPR (e. g. this study) | 1:1000 | (39) |
| rabbit anti-ISCU2 | Lill laboratory, validated in house by RNAi | 1:400 | (41) |
| rabbit anti-NBP35 | Lill laboratory, validated in house by RNAi | 1:666 (affinity purified) | (34) |
| rabbit anti-NDOR1 | Validated by RNAi and immunoprecipitation (this study and personal observation) | 1:666 (plus ABC) | Proteintech (11404-1-ap, RRID:AB2251225) |
| rabbit anti-NFS1 | Lill laboratory, validated in house by RNAi | 1:1000 | (35) |
| mouse anti-NFS1 (clone B-7) | validated in house by RNAi | 1:400 | Santa Cruz Technology (sc-365308, RRID:AB_10843245) |
| mouse anti-alpha tubulin (clone DM1α) | validated by the manufacturer and in house by immunofluorescence (personal observation) | up to 1:10,000 | Sigma-Aldrich (T9026, RRID:AB_477593) |

| **General** | | | |
|---|---|---|---|
| **Antibody** | **Validation** | **Dilution used** | **Source (product #)** |
| mouse anti-FLAG (clone M2) | Validated in house by immunoprecipitation (this study) | n. a. (affinity gel) | Sigma-Aldrich (A2220, RRID:AB_10063035) |
| rabbit anti-HA | Validated in house by immunoprecipitation using yeast cells transformed with plasmids lacking HA epitope tag, this study | 1:1000 | Santa Cruz (SC-7392 AC, RRID:AB_627809) |
| rabbit anti-myc | Validated in house by immunoprecipitation using yeast cells transformed with plasmids lacking myc epitope tag, this study | 1:1000 | Santa Cruz (SC-40 AC, RRID:AB_2857941) |
| Mouse anti-c-myc (clone 9E10) | Validated by the developer (JM Bishop; 10.1128/mcb.5.12.3610-3616.1985) | n. a. (affinity gel) | Developmental Studies Hybridoma Bank |

| | | | |
|---|---|---|---|
| | | | (deposited by JM Bishop) RRID:AB_ 10637885 |
| anti-TAP (IgG Sepharose) | Validated in house by immunoprecipitation using yeast cells transformed with plasmids lacking TAP epitope tag, this study | 1:1000 | Sigma-Aldrich (GE17-0969-01) |
| peroxidase-conjugated goat anti-rabbit Ig | Validated by manufacturer | 1:7500 | Biorad (170-6515, RRID:AB_1112 5142) |
| peroxidase-conjugated goat anti-mouse Ig | Validated by manufacturer | 1:7500 | Biorad (170-6516, RRID:AB_1112 5547) |
| biotin-conjugated goat anti-rabbit Ig | Validated by manufacturer | 1:2500 | Vector Laboratories (BA-1000, RRID:AB_2313 606) |
| Avidin-Biotin-Complex (ABC) system | Validated by manufacturer | n. a. | Vector Laboratories (PK-6100, RRID:AB_2336 819) |

### References to Supplement

1. Mühlenhoff U, *et al.* (2010) Cytosolic monothiol glutaredoxins function in intracellular iron sensing and trafficking via their bound iron-sulfur cluster. *Cell Metab* 12(4):373-385.
2. Pujol-Carrion N, Belli G, Herrero E, Nogues A, & de la Torre-Ruiz MA (2006) Glutaredoxins Grx3 and Grx4 regulate nuclear localisation of Aft1 and the oxidative stress response in Saccharomyces cerevisiae. *J Cell Sci* 119(Pt 21):4554-4564.
3. Li G, Nanjaraj Urs AN, Dancis A, & Zhang Y (2022) Genetic suppressors of Deltagrx3 Deltagrx4, lacking redundant multidomain monothiol yeast glutaredoxins, rescue growth and iron homeostasis. *Biosci Rep* 42(6).
4. Mercier A & Labbe S (2009) Both Php4 function and subcellular localization are regulated by iron via a multistep mechanism involving the glutaredoxin Grx4 and the exportin Crm1. *J Biol Chem* 284(30):20249-20262.
5. Misslinger M, *et al.* (2019) The monothiol glutaredoxin GrxD is essential for sensing iron starvation in Aspergillus fumigatus. *PLoS Genet* 15(9):e1008379.
6. Attarian R, *et al.* (2018) The Monothiol Glutaredoxin Grx4 Regulates Iron Homeostasis and Virulence in Cryptococcus neoformans. *mBio* 9(6).
7. Alkafeef SS, *et al.* (2020) Proteomic profiling of the monothiol glutaredoxin Grx3 reveals its global role in the regulation of iron dependent processes. *PLoS Genet* 16(6):e1008881.
8. Wang Z, *et al.* (2019) A fungal ABC transporter FgAtm1 regulates iron homeostasis via the transcription factor cascade FgAreA-HapX. *PLoS Pathog* 15(9):e1007791.
9. Cha H, *et al.* (2008) PICOT is a critical regulator of cardiac hypertrophy and cardiomyocyte contractility. *J Mol Cell Cardiol* 45(6):796-803.
10. Cheng NH, *et al.* (2011) A mammalian monothiol glutaredoxin, Grx3, is critical for cell cycle progression during embryogenesis. *FEBS J* 278(14):2525-2539.
11. Pham K, *et al.* (2017) Loss of glutaredoxin 3 impedes mammary lobuloalveolar development during pregnancy and lactation. *Am J Physiol Endocrinol Metab* 312(3):E136-E149.
12. Cheng N, *et al.* (2021) Crucial Role of Mammalian Glutaredoxin 3 in Cardiac Energy Metabolism in Diet-induced Obese Mice Revealed by Transcriptome Analysis. *Int J Biol Sci* 17(11):2871-2883.
13. Donelson J, *et al.* (2019) Cardiac-specific ablation of glutaredoxin 3 leads to cardiac hypertrophy and heart failure. *Physiol Rep* 7(8):e14071.
14. Cheng NH, *et al.* (2011) Arabidopsis monothiol glutaredoxin, AtGRXS17, is critical for temperature-dependent postembryonic growth and development via modulating auxin response. *J Biol Chem* 286(23):20398-20406.
15. Knuesting J, *et al.* (2015) Arabidopsis glutaredoxin S17 and its partner, the nuclear factor Y subunit C11/negative cofactor 2alpha, contribute to maintenance of the shoot apical meristem under long-day photoperiod. *Plant Physiol* 167(4):1643-1658.
16. Mortimer RK & Johnston JR (1986) Genealogy of principal strains of the yeast genetic stock center. *Genetics* 113(1):35-43.
17. Sikorski RS & Hieter P (1989) A system of shuttle vectors and yeast host strains designed for efficient manipulation of DNA in *Saccharomyces cerevisiae. Genetics* 122:19-27.
18. Sipos K, *et al.* (2002) Maturation of cytosolic iron-sulfur proteins requires glutathione. *J. Biol. Chem.* 277:26944-26949.
19. Stegmaier K, *et al.* (2019) Apd1 and Aim32 Are Prototypes of Bishistidinyl-Coordinated Non-Rieske [2Fe-2S] Proteins. *J Am Chem Soc* 141(14):5753-5765.
20. Mühlenhoff U, Gerber J, Richhardt N, & Lill R (2003) Components involved in assembly and dislocation of iron-sulfur clusters on the scaffold protein Isu1p. *EMBO J.* 22:4815-4825.
21. Schulz V, *et al.* (2022) Functional spectrum and specificity of mitochondrial ferredoxins FDX1 and FDX2. *Nat Chem Biol.*
22. Srinivasan V, Pierik AJ, & Lill R (2014) Crystal structures of nucleotide-free and glutathione-bound mitochondrial ABC transporter Atm1. *Science* 343(6175):1137-1140.
23. Janke C, *et al.* (2004) A versatile toolbox for PCR-based tagging of yeast genes: new fluorescent proteins, more markers and promoter substitution cassettes. *Yeast* 21(11):947-962.

24. Hausmann A, *et al.* (2005) The eukaryotic P-loop NTPase Nbp35: An essential component of the cytosolic and nuclear iron-sulfur protein assembly machinery. *Proc. Natl. Acad. Sci. USA* 102:3266-3271.
25. Balk J, Aguilar Netz DJ, Tepper K, Pierik AJ, & Lill R (2005) The essential WD40 protein Cia1 is involved in a late step of cytosolic and nuclear iron-sulfur protein assembly. *Mol Cell Biol* 25(24):10833-10841.
26. Balk J, Pierik AJ, Aguilar Netz D, Mühlenhoff U, & Lill R (2004) The hydrogenase-like Nar1p is essential for maturation of cytosolic and nuclear iron-sulphur proteins. *EMBO J.* 23:2105-2115.
27. Braymer JJ, Stumpfig M, Thelen S, Muhlenhoff U, & Lill R (2019) Depletion of thiol reducing capacity impairs cytosolic but not mitochondrial iron-sulfur protein assembly machineries. *Biochim Biophys Acta Mol Cell Res* 1866(2):240-251.
28. Rietzschel N, Pierik AJ, Bill E, Lill R, & Muhlenhoff U (2015) The basic leucine zipper stress response regulator Yap5 senses high-iron conditions by coordination of [2Fe-2S] clusters. *Molecular and cellular biology* 35(2):370-378.
29. Netz DJ, Pierik AJ, Stümpfig M, Mühlenhoff U, & Lill R (2007) The Cfd1-Nbp35 complex acts as a scaffold for iron-sulfur protein assembly in the yeast cytosol. *Nat Chem Biol* 3(5):278-286.
30. Netz DJ, *et al.* (2012) A bridging [4Fe-4S] cluster and nucleotide binding are essential for function of the Cfd1-Nbp35 complex as a scaffold in iron-sulfur protein maturation. *J Biol Chem* 287(15):12365-12378.
31. Netz DJ, *et al.* (2010) Tah18 transfers electrons to Dre2 in cytosolic iron-sulfur protein biogenesis. *Nat Chem Biol* 6(10):758-765.
32. Paul VD, *et al.* (2015) The deca-$GX_3$ proteins Yae1-Lto1 function as adaptors recruiting the ABC protein Rli1 for iron-sulfur cluster insertion. *eLife* 4:e08231.
33. Hausmann A, Samans B, Lill R, & Muhlenhoff U (2008) Cellular and Mitochondrial Remodeling upon Defects in Iron-Sulfur Protein Biogenesis. *J Biol Chem* 283(13):8318-8330.
34. Stehling O, *et al.* (2008) Human Nbp35 is essential for both cytosolic iron-sulfur protein assembly and iron homeostasis. *Molecular and cellular biology* 28(17):5517-5528.
35. Biederbick A, *et al.* (2006) Role of human mitochondrial Nfs1 in cytosolic iron-sulfur protein biogenesis and iron regulation. *Mol. Cell. Biol.* 26:5675-5687.
36. Ran FA, *et al.* (2013) Genome engineering using the CRISPR-Cas9 system. *Nat Protoc* 8(11):2281-2308.
37. Brummelkamp TR, Bernards R, & Agami R (2002) A system for stable expression of short interfering RNAs in mammalian cells. *Science* 296(5567):550-553.
38. Haunhorst P, *et al.* (2013) Crucial function of vertebrate glutaredoxin 3 (PICOT) in iron homeostasis and hemoglobin maturation. *Mol Biol Cell* 24(12):1895-1903.
39. Brautigam L, *et al.* (2011) Vertebrate-specific glutaredoxin is essential for brain development. *Proc Natl Acad Sci U S A* 108(51):20532-20537.
40. Stehling O, *et al.* (2018) Function and crystal structure of the dimeric P-loop ATPase CFD1 coordinating an exposed [4Fe-4S] cluster for transfer to apoproteins. *Proc Natl Acad Sci U S A* 115(39):E9085-E9094.
41. Navarro-Sastre A, *et al.* (2011) A fatal mitochondrial disease is associated with defective NFU1 function in the maturation of a subset of mitochondrial Fe-S proteins. *Am J Hum Genet* 89(5):656-667.
42. Upadhyay AS, *et al.* (2014) Viperin is an iron-sulfur protein that inhibits genome synthesis of tick-borne encephalitis virus via radical SAM domain activity. *Cell Microbiol* 16(6):834-848.
43. Doench JG, *et al.* (2016) Optimized sgRNA design to maximize activity and minimize off-target effects of CRISPR-Cas9. *Nat Biotechnol* 34(2):184-191.
44. Heigwer F, Kerr G, & Boutros M (2014) E-CRISP: fast CRISPR target site identification. *Nat Methods* 11(2):122-123.
45. Bekri S, *et al.* (2000) Human ABC7 transporter: Gene structure and mutation causing X-linked sideroblastic anemia with ataxia (XLSA/A) with disruption of cytosolic iron-sulfur protein maturation. *Blood* 96:3256-3264.
46. Zheng L, Baumann U, & Reymond JL (2004) An efficient one-step site-directed and site-saturation mutagenesis protocol. *Nucleic acids research* 32(14):e115.

47. Arakawa H, Lodygin D, & Buerstedde JM (2001) Mutant loxP vectors for selectable marker recycle and conditional knock-outs. *BMC Biotechnol* 1:7.
48. Stehling O, Paul VD, Bergmann J, Basu S, & Lill R (2018) Biochemical Analyses of Human Iron–Sulfur Protein Biogenesis and of Related Diseases. *Methods Enzymol.* 599:227-263.
49. Lill R & Freibert SA (2020) Mechanisms of Mitochondrial Iron-Sulfur Protein Biogenesis. *Annu Rev Biochem* 89:471-499.
